# ScReNI: single-cell regulatory network inference through integrating scRNA-seq and scATAC-seq data

**DOI:** 10.1101/2024.09.10.612385

**Authors:** Xueli Xu, Yanran Liang, Miaoxiu Tang, Jiongliang Wang, Xi Wang, Yixue Li, Jie Wang

## Abstract

Single cells exhibit heterogeneous gene expression profiles and chromatin accessibility, measurable separately via single-cell RNA sequencing (scRNA-seq) and single-cell transposase chromatin accessibility sequencing (scATAC-seq). Consequently, each cell possesses a unique gene regulatory network. However, limited methods exist for inferring cell-specific regulatory networks, particularly through the integration of scRNA-seq and scATAC-seq data. Here, we develop a novel algorithm named single-cell regulatory network inference (ScReNI), which leverages *k*-nearest neighbors and random forest algorithms to integrate scRNA-seq and scATAC-seq data for inferring gene regulatory networks at the single-cell level. ScReNI is built to analyze both paired and unpaired datasets for scRNA-seq and scATAC-seq. Using these two types of single-cell sequencing datasets, we validate that a higher fraction of regulatory relationships inferred by ScReNI are detected by chromatin immunoprecipitation sequencing (ChIP-seq) data. ScReNI shows superior performance in network-based cell clustering when compared to existing single-cell network inference methods. Importantly, ScReNI offers the unique function of identifying cell-enriched regulators based on each cell-specific network. In summary, ScReNI facilitates the inferences of cell-specific regulatory networks and cell-enriched regulators.

## INTRODUCTION

Transcriptional regulatory networks are essential for cells to determine which genes should be transcribed (1). Typically, *trans*-regulators (e.g., transcription factors) bind to *cis*-regulatory elements to regulate gene transcription. Gene transcription levels can be globally measured using RNA sequencing (RNA-seq) technology (2). The genomic locations of *cis*-regulatory elements, which are commonly present in open chromatin regions, can be detected using the assay for transposase-accessible chromatin with sequencing (ATAC-Seq) (3). The revolution of single-cell sequencing technology has led to the advent of single-cell RNA-seq (scRNA-seq) and single-cell ATAC-seq (scATAC-seq) (4). These cutting-edge technologies enable the measurement of gene expression and chromatin accessibility at the single-cell level, either in different batches of cells or simultaneously within the same cells, resulting in either unpaired or paired datasets for scRNA-seq and scATAC-seq, respectively (5). These single-cell sequencing data offer unprecedented opportunities for the accurate inference of regulatory networks.

Regulatory network analysis has become an indispensable tool for uncovering the underlying regulatory mechanisms and key regulatory factors that govern a myriad of biological processes (6–8). By leveraging bulk or single-cell RNA-seq data, algorithms such as GENIE3, PIDC, and SCENIC, have been employed to infer cell type-specific regulatory networks (9–11). Several methods like SCENIC+ and IReNA emerged, offering enhanced capabilities to infer cell type-specific regulatory networks by integrating scRNA-seq and scATAC-seq data (12, 13). It is noteworthy that while cells from the same cell type may share similar regulatory networks, each cell exhibits a unique regulatory network due to its heterogeneous gene expression profile and chromatin accessibility. This uniqueness highlights the importance of inferring cell-specific regulatory networks, which are essential for uncovering the regulatory heterogeneity and key regulators at the single-cell level.

Currently, only few methods have been developed to infer cell-specific networks, primarily tailored for scRNA-seq data. For instance, CSN (cell-specific network) is designed to infer networks for each individual cell using scRNA-seq data (14). It calculates binary values through a statistical measurement to quantify the dependencies between any two genes, thereby indicating gene regulatory relationships. LIONESS (linear interpolation to obtain network estimates for single samples) has been used to infer networks through extracting sample-specific regulatory networks from population-level networks (15). Furthermore, CeSpGRN (cell specific gene regulatory network inference) reconstructs cell-specific regulatory networks from scRNA-seq data. When paired scATAC-seq data is available, CeSpGRN only uses chromatin accessiblity to narrow down the regulatory relationships inferred from scRNA-seq data, but does not establish the essential relationships between scRNA-seq and scATAC-seq (16). The method of integrating scRNA-seq and scATAC-seq data to infer cell-specific regulatory networks is still quite limited, particularly when dealing with unpaired scRNA-seq and scATAC-seq data.

In this study, we develop a new computational method named ScReNI, which constructs cell-specific regulatory networks from scRNA-seq and scATAC-seq data in an unpaired or paired manner. Specifically, ScReNI initially integrates unpaired scRNA-seq and scATAC-seq datasets through aligning them in a shared analytical space. It then establishes the association between genes and peaks across all cells. Finally, ScReNI uses *k*-nearest neighbors and random forest algorithms to infer gene regulatory relationships for individual cells by modeling the integrated scRNA-seq and scATAC-seq data. To assess the performance of ScReNI compared to existing methods, we also utilize scRNA-seq data alone to infer cell-specific networks. Through validation using unpaired and paired single-cell sequencing datasets, we demonstrate that a higher fraction of regulatory relationships inferred by ScReNI are detected by chromatin immunoprecipitation sequencing (ChIP-seq) data. ScReNI shows superior performance in network-based cell clustering when compared to existing single-cell network inference methods. Importantly, ScReNI enables the identification of cell-enriched regulators based on cell-specific regulatory networks.

## MATERIAL AND METHODS

### Algorithm overview

ScReNI is developed to infer cell-specific regulatory networks from unpaired (or paired) scRNA-seq and scATAC-seq data (Figure 1 and Figure S1A). The count matrices of gene expression and chromatin accessibility are independently generated from scRNA-seq and scATAC-seq data, respectively. These matrices are then used to predict cell-specific regulatory networks through four key steps: clustering of cells measured by scRNA-seq and scATAC-seq, identification of *k*-nearest neighbors for each cell, inference of gene regulatory relationships via the random forest, and reconstruction and evaluation of cell-specific regulatory networks. After performing these four steps, regulators enriched in each cell will be identified through statistically analyzing cell-specific regulatory networks.

**Figure 1.**
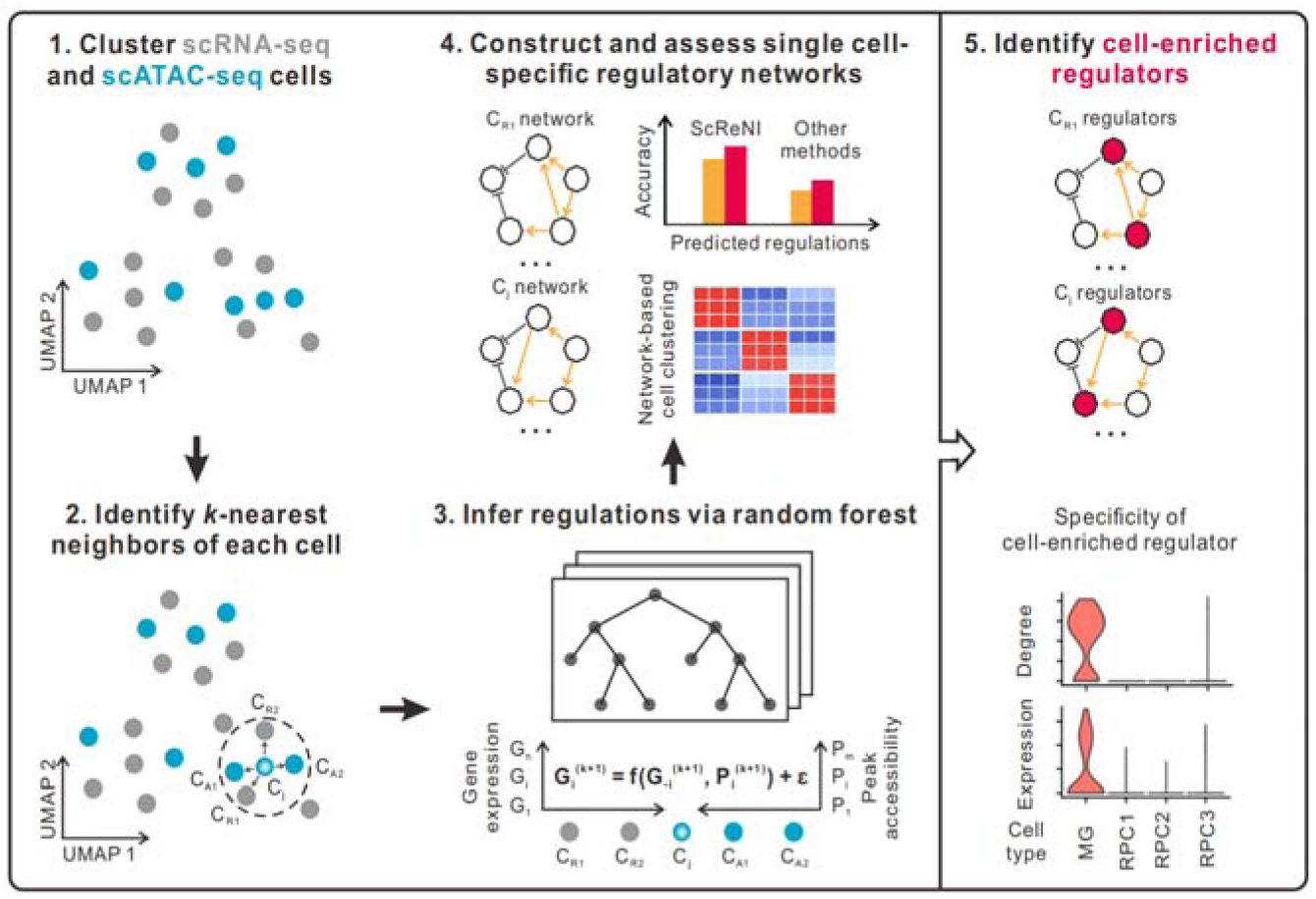
Overview of ScReNI. ScReNI contains five major parts. First, cells measured by scRNA-seq and scATAC-seq are clustered together. Second, neighbors for each cell are identified using the *k*-nearest neighbors algorithm. Third, the weights of transcription factors regulating target genes are calculated using random forest. Fourth, ScReNI is validated by assessing cell-specific regulatory networks. Fifth, cell-enriched regulators are identified based on each cell-specific network.

In ScReNI, we employ the weighted-nearest neighbor analysis from Seurat software to integrate paired scRNA-seq and scATAC-seq data (17). This analytical strategy allows us to identify the nearest neighbors for each cell and calculate the relative utility of each data type from scRNA-seq and scATAC-seq. In contrast, for the integration of unpaired scRNA-seq and scATAC-seq data, we apply the Seurat anchoring procedure to identify anchors between the two data types, followed by cell clustering using Harmony (18, 19). Subsequently, we identify neighbor cells of each cell based on single-cell clustering and the weighted-nearest neighbors algorithm. The cell itself and its neighbor cells are utilized to infer the regulatory network of the single cell.

The determination of associations between genes and peaks in ScReNI contains a two-step process (Figure S1B). First, genes are associated with peaks which are located within 250 kilobase pairs (kbp) upstream and 250 kbp downstream of transcription start sites of the gene. This restriction is strategically implemented to minimize the risk of generating false predictions. Using the ‘findOverlaps’ function from the R package ‘GenomicRanges’, we ascertained which genes overlapped with region-specific peaks. Second, the correlation between genes and peaks is established by calculating an absolute value of the Spearman correlation coefficient. Then, the ‘matchMotifs’ function from the R package ‘motifmatchr’ is employed to pinpoint potential binding sites of transcription factors within the genomic regions. Genes are associated with peaks if they are greater than 0.1 correlation.

ScReNI applies gene-peak associations to a group of cells comprising the cell of interest and its *k*-nearest neighbors. Utilizing random forest, ScReNI builds regression models with scRNA-seq and scATAC-seq data, thereby calculating the regulatory weights of transcription factors on their target genes. For each gene, the algorithm evaluates the influence of all other genes, including peak-associated transcription factors. The weights are then applied to reconstruct the regulatory network for an individual cell. This methodology, called weighted-nearest neighbor ScReNI (wScReNI), is primarily utilized for analyzing scRNA-seq and scATAC-seq data. The above method can also be used to analyze scRNA-seq data alone for cell-specific network inference, named *k*-nearest neighbor ScReNI (kScReNI), using the *k*-nearest neighbors algorithm to identify neighbor cells of each cell.

To benchmark the performance of ScReNI against existing methods, we reconstructed cell-specific networks using CSN, LIONESS, and CeSpGRN. These methods infer cell-specific regulatory networks solely based on transcriptomic data. For this purpose, we employed two types of publicly available datasets: one consisting of unpaired scRNA-seq and scATAC-seq data from retinal development, and the other containing paired scRNA-seq and scATAC-seq data from peripheral blood mononuclear cells of a healthy donor. We utilized these datasets to validate the inferred cell-specific regulatory networks. Subsequently, downstream analyses were conducted, which included the identification of regulators specifically enriched in individual cells, based on the reconstructed cell-specific regulatory networks.

### Datasets used for ScReNI

Two types of high-quality scRNA-seq and scATAC-seq data were measured separately in unpaired and paired manners to assess the performance of our ScReNI method.

The unpaired scRNA-seq and scATAC-seq data were publicly available from the study of retinal development in mice (20). The original research reported thirteen major retinal cell types. For our analysis, we focused on three subtypes of retinal progenitor cells (RPCs), designated as RPC1, RPC2, and RPC3, as well as Müller glial (MG) cells, utilizing their original cell type annotations.

The paired scRNA-seq and scATAC-seq data were obtained from peripheral blood mononuclear cells (PBMCs) of a healthy donor which was obtained by 10x Genomics. The libraries for paired ATAC and gene expression were generated from the isolated nuclei as described in the Chromium Next GEM Single Cell Multiome ATAC + Gene Expression User Guide (CG000338 Rev A) and sequenced on Illumina Novaseq 6000 v1 Kit (Forward Strand Dual-Index Workflow). For our analysis, we selected single-cell sequencing data from CD14 monocytes, CD16 monocytes, CD4 naive cells, CD8 naive cells, conventional dendritic cells (cDC) cells, memory B cells, natural killer (NK) cells, and regulatory T cells (Treg) cells.

### Integration of scRNA-seq and scATAC-seq data

We initiate our analysis by generating an estimate of the transcriptional activity of each gene. This is accomplished by quantifying ATAC-seq counts within the 2 kb-upstream region and gene body, using the GeneActivity function in the Signac package (21). The resulting gene activity scores, derived from the scATAC-seq data, serve as inputs for canonical correlation analysis. Following this, we apply the Seurat v5 anchoring procedure to identify anchors between the scRNA-seq and scATAC-seq data (22). Once anchors are identified, we facilitate the transfer of annotations from the scRNA-seq dataset onto the scATAC-seq cells. Following the anchoring, we proceed with cell clustering using Harmony (19).

### Identification of the nearest neighbors for each cell

For each cell, the weighted nearest neighbor procedure implemented in R package Seurat v5 learns a set of modality weights (17). The identification of the nearest neighbors of the single cell is achieved through a multi-step process. Initially, a Seurat object is constructed from scRNA-seq data. Subsequently, this is complemented by integrating scATAC-seq data as an additional assay within the same Seurat object. Both assays are then independently subjected to standard pre-processing and dimensionality reduction which are specific to data types. The culmination of these steps is the computation of a weighted nearest neighbor graph, which effectively combines the information from both scRNA-seq and scATAC-seq modalities. This graph is essential for subsequent uniform manifold approximation and projection (UMAP) visualization and clustering, providing a comprehensive framework for identifying the nearest neighbors.

### Regulatory network inference in ScReNI

ScReNI is a new tool designed to analyze scRNA-seq and scATAC-seq data, accommodating both unpaired and paired datasets. For unpaired datasets, ScReNI initiates the analysis with cell clustering using Seurat anchor-based integration and Harmony. ScReNI operates on a dataset comprising *N* samples (cells) and *G+ P* features (*G* genes and *P* peaks).

The main hypothesis is that the expression of a target gene is influenced by the transcription factors and the peaks associated with the target gene. Let *x^-q^* represent the vector of expression values for all genes except gene *q*, represented as

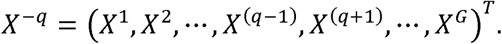

We propose that the expression level *x^q^* of the target gene *q* is regulated by the transcription factors present in *x^-q^* and the chromatin-accessible peaks in *Y^q^*. This regulatory relationship is formalized in the model:

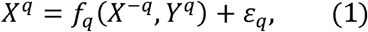

where *ε_q_* represents random noise with a mean of zero, and the function *f_q_* encapsulates the regulatory effects of the transcription factors in *X^-q^* on gene *q*, considering a subset of the peaks in *Y^q^* that are associated with gene *q*.

ScReNI predicts regulatory links pointing to gene *q* by learning the function *f_q_*. Within the ScReNI framework, the regulatory function *f_q_* is modeled using a random forest method, employing the default parameters. The random forest method returns the important scores of all transcription factors in *x^-q^* and peaks in *Y^q^* using IncNodePurity and IncMSE metrics.

### Determination of gene regulatory relationships

The weight *w_i,j_* between gene *i* and *J* is determined by aggregating the importance scores of gene *J* and the associated peaks of gene *i*. This metric quantifies the regulatory strength of gene *J* on gene *i*, integrating the information from chromatin accessibility data pertinent to gene *i*. The weight *w_i,j_* between gene *i* and *J* is calculated using the following formula:

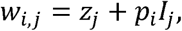

where *z_j_* denotes the importance score of gene *j*, *p_i_* represents the importance score of peaks associated with gene *i*, and *l_j_* refers to an indicator of whether gene *J* is related to the peaks associated with gene *i*. There are three scenarios to calculate the weight:

1. If gene *j* (where *i* and *j* are different genes) is not a transcription factor, the weight *w_i,j_* between gene *i* and gene *j* is equivalent to the importance score of gene *j*.
2. If gene *j* is a transcription factor but does not have an association with peak *i*, the regulatory strength of gene *j* on gene *i* is determined by the importance score of gene *j* alone.
3. If gene *j* is a transcription factor and has an association with peak *i*, the regulatory strength of gene *j* on gene *i* is the sum of the importance scores of both gene *j* and peak *i*.

### Construction of cell-specific regulatory networks

For a given single cell *c* (*c* = 1, 2, ·, *N*), the *k* nearest neighbor cells are identified through a weighted nearest neighbor analysis. Cell *c* and its neighboring cells are then used as inputs of equation (1) to infer the regulatory network of the single cell *c*. The obtained cell-specific regulatory networks are directed from regulators to targeted genes. The weights between genes represented the regulatory strengths.

### Calculation of network similarity

For each gene regulatory network, regulation pairs are ranked according to their weights. Based on the top 3,000 regulation pairs, we calculate the number of regulatory relationships shared by two different cells to measure the similarity between the corresponding networks of the cells.

### Baseline models

CSN, LIONESS, and CeSpGRN were used as the baseline models to infer cell-specific regulatory networks (14–16). All default settings were used.

1. CSN analyzes scRNA-seq data at the single-cell level. It employs a statistical model to identify gene-gene associations based on their joint and marginal probabilities.
2. LIONESS estimates individual-specific regulatory networks from gene expression data. The method is based on the principle that the presence or absence of a single sample can cause a slight change in an overall network model. This change is utilized to assess the impact of a sample on the aggregate network, which in turn helps to determine a sample-specific network.
3. CeSpGRN employs a Gaussian weighted kernel to model the gene expression from scRNA-seq data and learn the gene regulatory network of a given cell. The kernel is constructed from the similarity of gene expressions or spatial locations between cells. When paired scATAC-seq data is available, CeSpGRN connects the TFs to their corresponding target genes based on the chromatin accessibility measured by scATAC-seq, and then narrows down the regulatory relationships between TFs and target genes inferred from the scRNA-seq data.

### Evaluation of the precision and recall on predicting regulatory networks

ChIP-Atlas was used to assess the inferred regulatory relationships (23, 24). We introduce the ChIP-Atlas dataset, a widely used gold standard network dataset that contains regulatory interactions detected by high-throughput sequencing experiments, such as ChIP-seq and ATAC-seq. This dataset serves as a valuable reference for evaluating the performance of different network inference methods. Precision and recall serve as metrics for evaluating the performance of different single-cell regulatory network inference methods. The results are visualized using histograms or boxplots for each single-cell regulatory network inference method. The thresholds are chosen to prioritize gene pairs with a different number of top regulation pairs which are ranked based on regulation weights.

### Clustering of single cells based on cell-specific networks

For network-based cell clustering, we calculated gene degrees derived from cell-specific networks. To evaluate the influence of the number of pairwise relationships between TFs and target genes, we applied diverse thresholds to the ranked weights. Specifically, we select the top 500, 800, 1000, and 3000 gene regulation pairs. To quantitatively measure the similarity between the resulting clustering and the known ground truth of cell types, we employ the adjusted rand index (ARI) as in the previous study (25). This metric is implemented in the ‘randIndex’ function from the R package ‘flexclust’, providing a robust statistical measure for evaluating the accuracy of our clustering method (26).

### Identification of cell-enriched regulators

To investigate pseudotime trajectories in scRNA-seq data of mouse retinal development, we transferred the data to Monocle3 using SeuratWrappers (27). Cells were clustered via UMAP reduction, and the principal graph was determined using the learn_graph function, with parameters set to euclidean_distance_ratio = 5, geodesic_distance_ratio = 2, and minimal_branch_len = 1000. Other parameters were kept at their default settings. Any RPC1 cell was set as the root when running order_cells. These data were then added back to Seurat as metadata for further analysis.

Considering the sparsity of scRNA-seq data, we employed the smoothed expression profiles for gene co-expression analysis. Pseudotime was divided into 50 equal intervals. Highly variable genes and expressed transcription factors were grouped into distinct modules using K-means clustering on the smoothed expression profiles. The optimal number of modules was determined by calculating the average silhouette score for K-means clustering, using the R package ‘cluster’. For each module, we conducted functional enrichment analysis using ClusterProfiler, leveraging Gene Ontology (GO) and Kyoto Encyclopedia of Genes and Genomes (KEGG) databases (28).

Based on modules of genes, the inferred regulatory network from each cell was modularized. The hypergeometric test with false discovery rate multiple hypotheses test correction was applied to calculate and calibrate the probability *P* that a given transcription factor is associated with the regulation of a specific gene module. Transcription factors identified as significantly enriched regulators of gene modules were determined using a threshold of 0.01 false discovery rate.

### Statistical analysis

All statistical analyses were conducted using R (v.4.3.3) software. Spearman rank correlation coefficients were used to examine the association between network degrees and expression levels of each cell-enriched regulator across all four cell types. Kruskal-Wallis test with false discovery rate correction was used to compare the statistical differences in network degrees, expression levels, and regulatory activities of cell-enriched regulators among four cell types (MG, RPC1, RPC2, and RPC3). The t-test with two-tailed was performed to statistically compare the mean number of transcription factors negatively and positively regulating the targeted gene. Statistical significance was set at p-value < 0.05.

## RESULTS

### ScReNI constructs cell-specific regulatory networks using unpaired scRNA-seq and scATAC-seq data

The unpaired scRNA-seq and scATAC-seq data were obtained from the previous study on mouse retinal development (20). We used scRNA-seq data and scATAC-seq data from age-matched nine time points between two sequencing types. The original study reported thirteen major retinal cell types. To illustrate ScReNI, we only included four cell types (RPC1, RPC2, RPC3, and MG). The original annotation was used as the growth truth in downstream analysis.

The scRNA-seq dataset encompassed 7,853 RPC1, 16,645 RPC2, 22,943 RPC3, and 936 MG cells, complemented by the scATAC-seq dataset with 6,049 RPC1, 10,464 RPC2, 11,912 RPC3, and 768 MG cells. Integration of these datasets was executed using Seurat anchor-based integration and Harmony (17, 19). For the single-cell clustering, we used the top 2000 highly variable genes and the top 10,000 highly variable peaks. The integrative analysis of scRNA-seq and scATAC-seq data showed a substantial overlap between the same cell types identified by two sequencing types (Figure 2A). The clustering results also delineated the developmental trajectory of MGs from RPCs, which was consistent with the original report (20). According to the cell clustering, we determined the neighboring cells for each cell using the weighted-nearest neighbors algorithm. Subsequently, we randomly chose 100 cells from each of RPCs and MGs as a representative subset of 400 cells for cell-specific network reconstruction. For each cell and its neighbors, gene regulatory networks were inferred using wScReNI. Then, the regulation pairs in regulatory networks were ranked according to the weights computed by wScReNI. Based on the top 3000 regulation pairs, we analyzed the similarities of gene regulatory networks separately within cell types and between cell types (Figure 2B). The median network similarities within the MG, PRC1, PRC2, and PRC3 were 282, 242, 205, and 203, respectively. The median network similarity between PRC1 and PRC2 was 192 which is the highest among all pairs of different cell types. The results indicated that each cell contained a specific and distinct regulatory network. Cells from the same cell type had more similar regulatory networks than those from different cell types.

**Figure 2.**
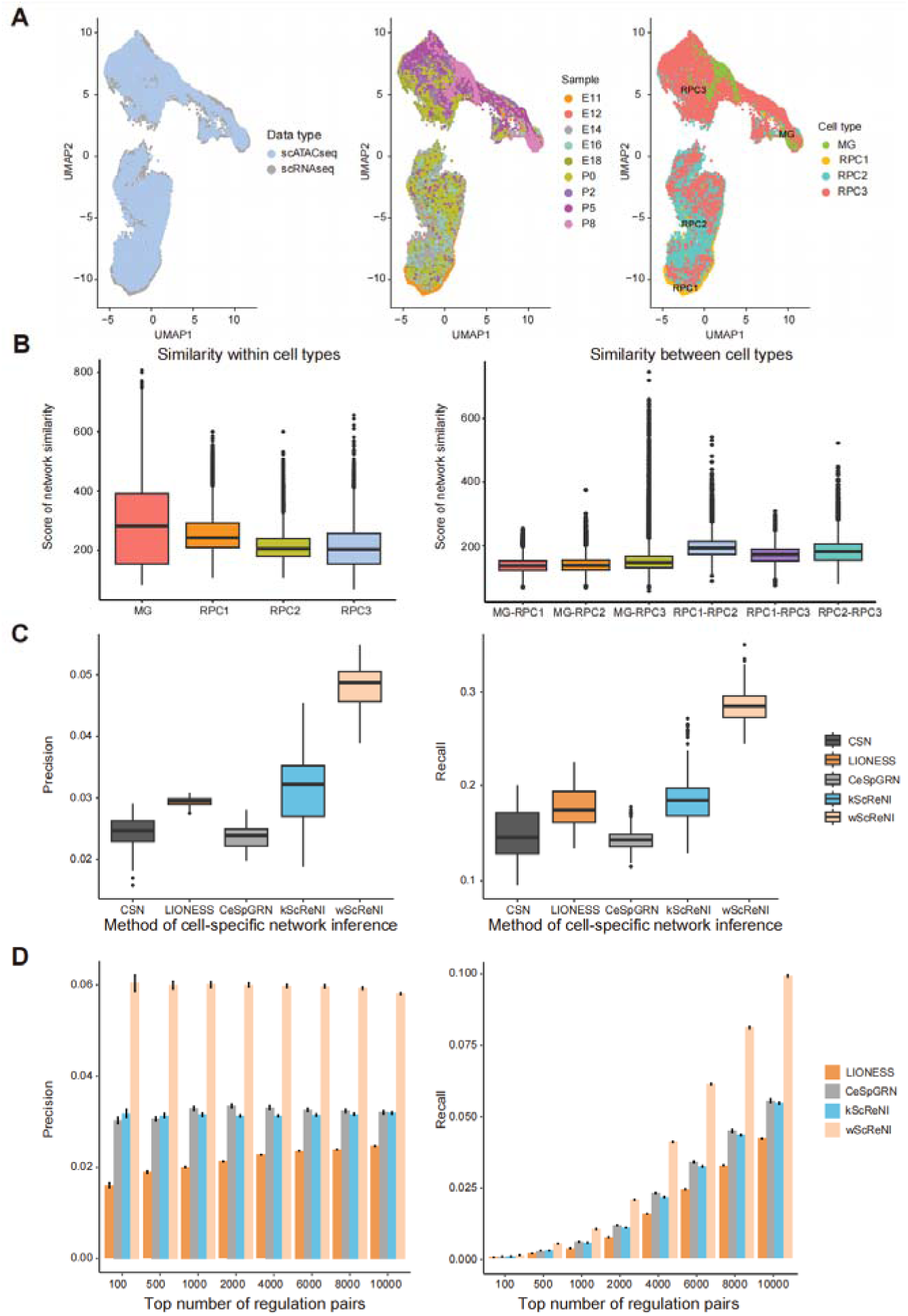
Cell-specific network analysis of unpaired scRNA-seq and scATAC-seq data from retinal development. (A) UMAP visualization of different data types, samples and cell types following multi-omics integration. (B) Similarities of gene regulatory networks separately within cell types and between cell types. (C) Precision and recall of single-cell regulatory relationships for different network inference methods using the same number of regulation pairs in CSN. (D) Precision and recall of single-cell regulatory relationships for four inference methods using different top numbers of regulation pairs.

### ScReNI shows superior performance on regulatory network inference

To assess the accuracy and reliability of the inferred regulatory relationships, we utilized the binding sites of transcription factors measured by ChIP-seq as the gold standard for gene regulatory relationships. The ChIP-seq data were from the ChIP-Atlas database (29). Following this, we calculated the precision and recall of cell-specific regulatory networks inferred by ScReNI, CSN, LIONESS, and CeSpGRN.

We first used the top 500 highly variable genes and the top 10,000 highly variable peaks for network inference. In network inference, wScReNI, kScReNI, LIONESS, and CeSpGRN calculated the continuous values for regulatory weights. However, the CSN method produced binary values for regulatory weights representing whether the gene is regulated or not. For each cell-specific regulatory network from retinal development, the regulation pairs were ranked based on their weights. To perform a fair comparison across different cell-specific network inference methods, we selected the same number of regulation pairs predicted by CSN. The results showed that both wScReNI and kScReNI had higher precision and recall than other methods (Figure 2C). The wScReNI showed the best performance among all methods. Specifically, the mean precision values for CSN, LIONESS, CeSpGRN, kScReNI, and wScReNI were 0.024, 0.029, 0.023, 0.031, and 0.047, respectively. The mean recall values for CSN, LIONESS, CeSpGRN, kScReNI, and wScReNI were 0.149, 0.179, 0.144, 0.185, and 0.288, respectively. We also calculated the precision and recall values of wScReNI, kScReNI, CeSpGRN, and LIONESS using different top numbers of regulation pairs (Figure 2D). Remarkably, wScReNI consistently outperformed other methods across all groups of cell-specific regulatory networks.

To confirm the above results, we also used the top 2000 highly variable genes and the top 10,000 highly variable peaks to infer cell-specific regulatory networks. The best performance was also observed in wScReNI regardless of the number of regulation pairs used in network inference (Figure S2A and S2B). These results indicated that ScReNI, especially wScReNI, had a remarkable performance on the accurate inference of cell-specific regulatory networks.

### ScReNI outperforms current methods in network-based cell clustering

To perform cell clustering analysis based on cell-specific networks, we computed the outdegrees of genes in each cell-specific network and obtained the matrix of gene outdegrees across all cells. According to gene degrees from the top 500 regulation pairs, single cells were clustered through Seurat and visualized using UMAP. The clustering based on gene degrees of cell-specific networks from wScReNI showed the best separation of different cell types (Figure 3A). We calculated the ARI to measure the similarity between the originally reported cell types and those assigned from network-based clustering. The results indicated that the wScReNI and kScReNI had higher ARI values than other methods. Specifically, the ARI values for CSN, LIONESS, CeSpGRN, kScReNI, and wScReNI were 0.488, 0.001, 0.513, 0.517, and 0.642, respectively. Then, we performed a hierarchical clustering and drew the heatmap using the top 500 regulation pairs. In wScReNI, the same cell types were more distinctly clustered together (Figure 3B). Notably, wScReNI and kScReNI exhibited higher ARI values compared to other methods. Specifically, the ARI values for CSN, LIONESS, CeSpGRN, kScReNI, and wScReNI were 0.226, 0.002, 0.238, 0.388, and 0.481, respectively. Network-based clustering analysis also indicated that ScReNI had better performance in distinguishing different cell types, even sub-cell types of RPCs (RPC1, RPC2, and RPC3). To further validate the performance, we conducted single-cell clustering based on different top regulation pairs, such as 500, 800, 1000, and 3000 regulation pairs. The results consistently showed that ScReNI outperformed the other methods(Table S1). Additionally, we also conducted network-based single-cell clustering using 2000 highly variable genes and different numbers of top regulation pairs. A similar result was observed that wScReNI showed the best performance overall, except for a few cases that were comparable to those obtained by the CeSpGRN method (Table S1).

**Figure 3.**
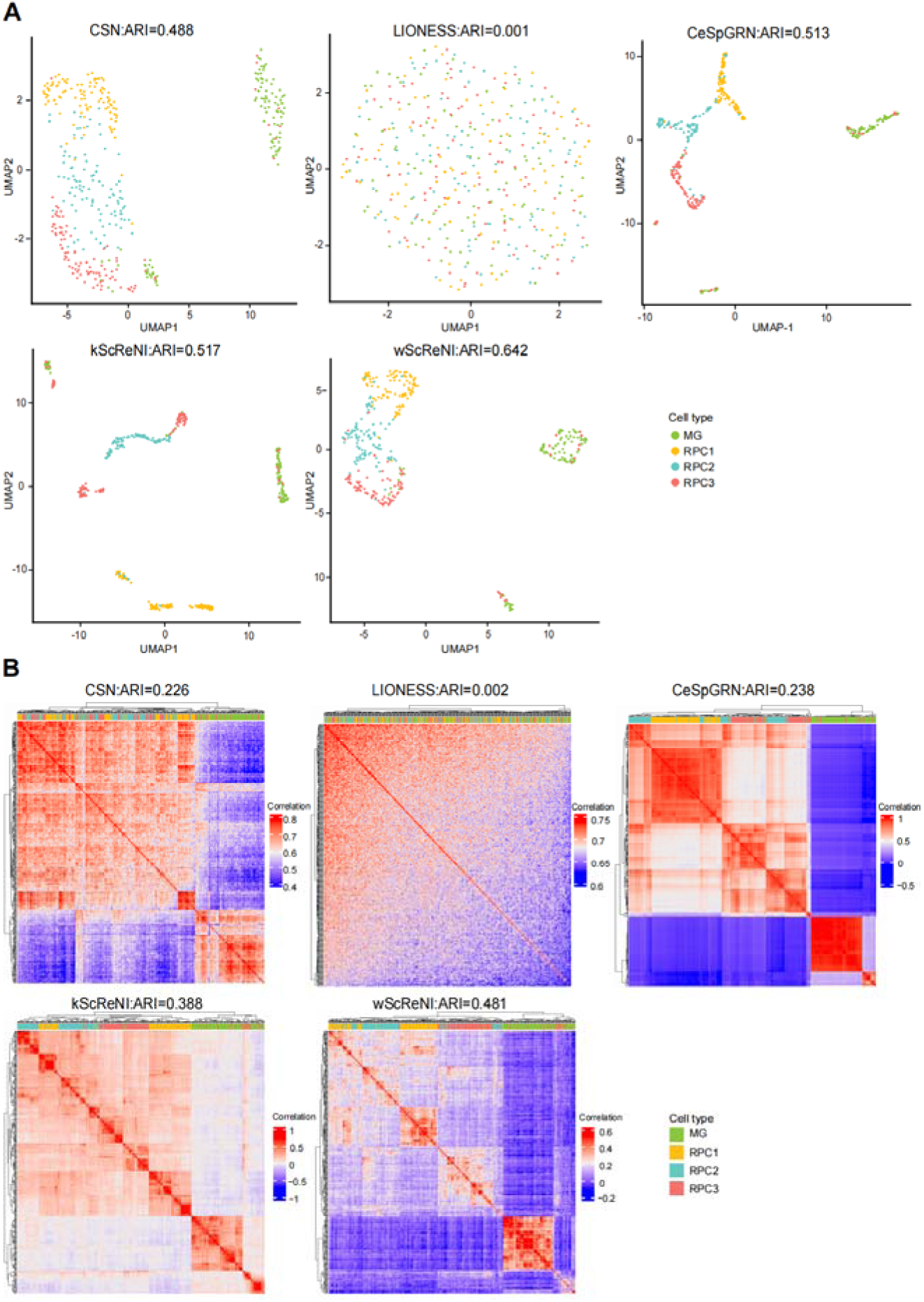
Network-based cell clustering of unpaired scRNA-seq and scATAC-seq data from retinal development. (A) UMAP of single cells using top 500 regulation pairs. (B) Hierarchical clustering of single cells using top 500 regulation pairs.

### ScReNI demonstrates superior performance in network inference using paired scRNA-seq and scATAC-seq data

To demonstrate the utility of ScReNI with paired scRNA-seq and scATAC-seq data, we employed publicly accessible single-cell multimodal datasets derived from peripheral blood mononuclear cells of a healthy donor, generated using the 10X Genomics platform. The dataset contains eight cell types, including 2812 CD14 monocytes, 514 CD16 monocytes, 1419 CD4 naive cells, 1410 CD8 naive cells, 198 cDC cells, 371 memory B cells, 468 NK cells, and 162 Treg cells. To construct cell-specific regulatory networks, we selected a representative subset of 400 cells, ensuring that each of the eight cell types contained 50 cells at random. The PBMC data were analyzed based on the top 500 highly variable genes and the top 10,000 highly variable peaks. Employing the weighted nearest-neighbor method, we performed multi-omics data clustering to establish a weighted combination graph of RNA and ATAC similarities. According to the single-cell clustering, we identified neighboring cells and inferred the regulatory network for each cell.

Using the same number of regulation pairs predicted by CSN, wScReNI showed significantly higher precision and recall compared to other methods (Figure 4A). The mean precision values for CSN, LIONESS, CeSpGRN, kScReNI, and wScReNI were 0.016, 0.016, 0.014, 0.014, and 0.025, respectively. The mean recall values for CSN, LIONESS, CeSpGRN, kScReNI, and wScReNI were 0.198, 0.193, 0.175, 0.177, and 0.300, respectively. Using various top numbers of regulation pairs for 500 highly variable genes, wScReNI consistently showed the highest precisions and recalls (Figure S3A). When 2000 highly variable genes were used, similar performance was observed (Figure S3B and S3C). These findings underscored the superior performance of wScReNI in identifying reliable regulatory relationships.

**Figure 4.**
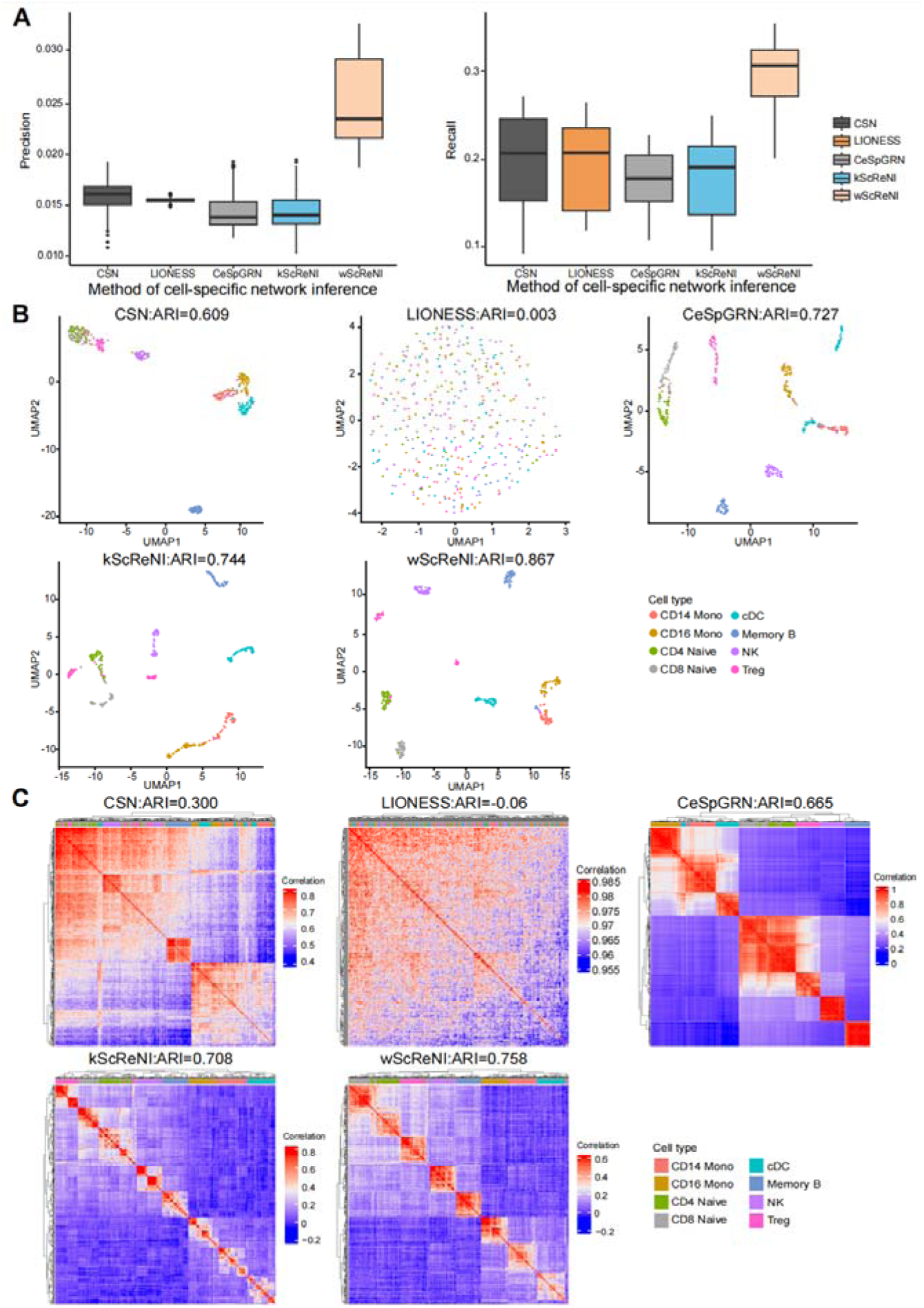
Network analysis of paired scRNA-seq and scATAC-seq data from PBMC. (A) Precision and recall of single-cell regulatory relationships using the same number of regulation pairs in CSN. (B) UMAP of single cells using top 500 regulation pairs. (C) Hierarchical clustering of single cells using top 500 regulation pairs.

In the analysis of the PBMC dataset, network-based cell clustering was also applied to the gene outdegrees of the top 500 regulatory pairs. UMAPs clearly illustrated that wScReNI provided better separation between different cell types compared to other methods (Figure 4B). This distinction was further corroborated by hierarchical clustering of the same gene degree matrix, which demonstrated distinct clustering patterns for various cell types within the wScReNI framework (Figure 4C). The ARI values were the highest in wScReNI, following kScReNI. Specifically, the ARI values from wScReNI were 0.867 and 0.758 for UMAPs and heatmaps, respectively. The results were highly consistent between the single-cell clustering and the ground truth of cell labels. The ARI values from CSN, LIONESS, CeSpGRN, and kScReNI were separately 0.609, 0.003, 0.727, and 0.744 in UMAPs. In heatmaps, kScReNI yielded an ARI value of 0.708, whereas CSN, LIONESS, and CeSpGRN were found to have ARI values not exceeding 0.670. We also chose the different top numbers of regulation pairs (500, 800, 1000, and 3000) to calculate gene degrees and cluster single cells. The ARI values provided strong evidence that wScReNI had superior performance against other methods in single-cell clustering analysis (Table S2). Additionally, we also conducted single-cell clustering based on 2000 highly variable genes and different top regulation pairs. Similar results were observed that wScReNI presented a better performance (Table S2). The consistent performance across different datasets indicated the robustness and reliability of wScReNI for regulatory network inference.

### ScReNI is applied to identify cell-enriched regulators

Using ScReNI, we sought to identify key regulators enriched in each cell-specific regulatory network, represented as cell-enriched regulators. To achieve this enrichment analysis, we performed a two-step analytical process inspired by IReNA (12). The first step is the division of all genes in cell-specific regulatory networks into different modules by applying K-means clustering to gene expression profiles across all cells. After genes in the cell-specific regulatory network were modularized, a hypergeometric test was performed to identify which factors significantly regulate each module of genes.

Cell-specific regulatory networks from retinal development inferred by wScReNI were utilized to illustrate the identification of cell-enriched regulators. To obtain reliable gene modules, we generated the smoothed gene expression profiles according to the pseudotime of single cells. The pseudotime of single cells was calculated based on the cell trajectory inferred by Monocle3 (27). We observed that the pseudotime was increased by the order of RPC1, RPC2, PRC3 and MG (Figure 5A). Based on the smoothed gene expression profiles, we clustered the top 2000 highly variable genes into six distinct modules (Figure 5B). Each gene module was associated with specific biological functions (Figure S4A). Specifically, the first module was related to organ development and cell proliferation. The second and third modules were involved in ribosome biogenesis and chromosome segregation, respectively. The genes in the other three modules were separately enriched in gliogenesis, visual perception, and synapse assembly. These enriched functions were closely associated with retinal development.

**Figure 5.**
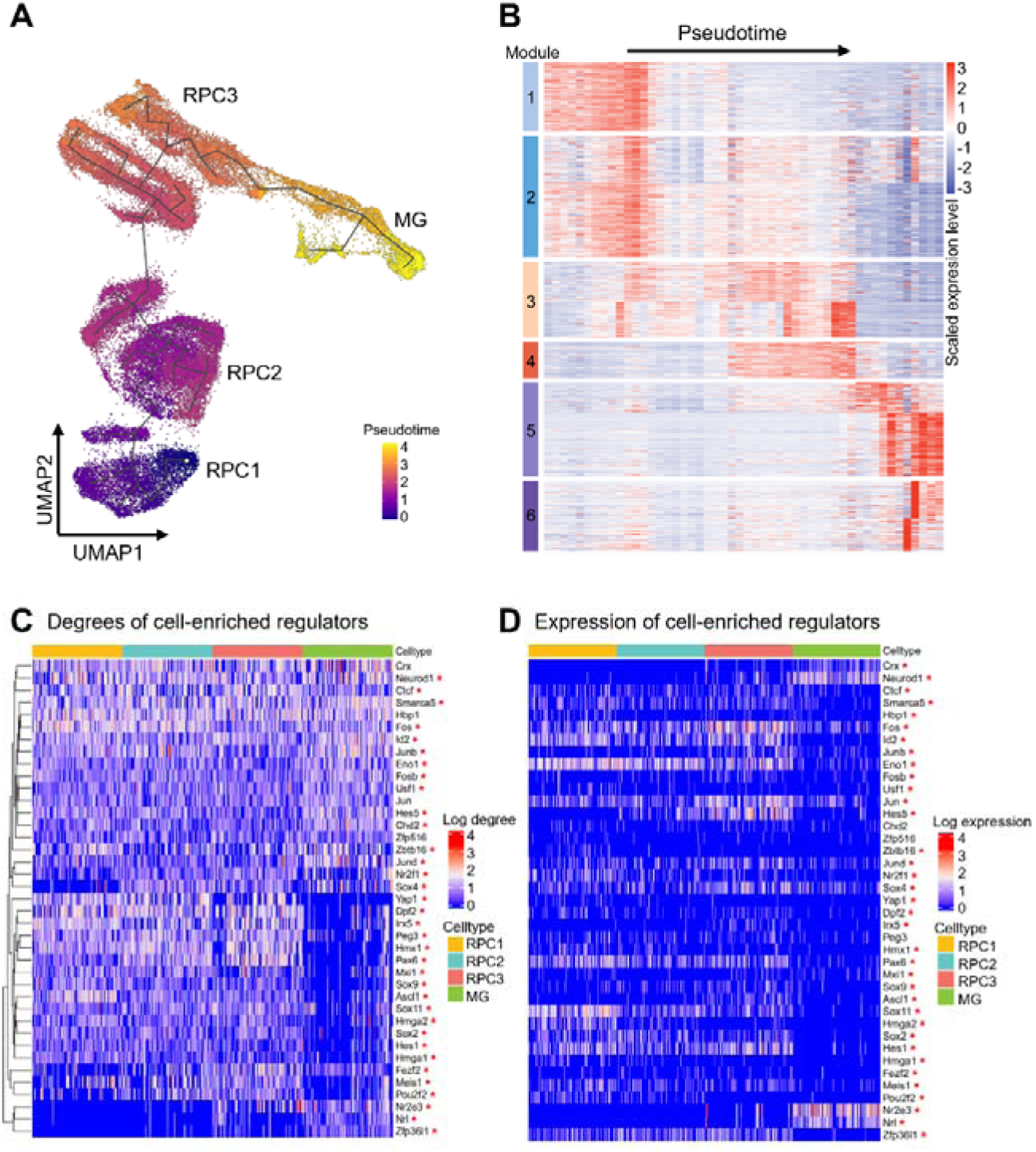
Identification of cell-enriched regulators during retinal development. (A) Pseudotime of single cells calculated by Monocle3. (B) Heatmap of genes divided by K-means. The color bars on the left represent six modules. (C) Degrees of cell-enriched regulators. (D) Expressions of cell-enriched regulators. P < 0.05 among four cell types was marked with a red star.

ScReNI successfully identified cell-enriched regulators for each modularized cell-specific regulatory network. For each cell type, we calculated the frequency of enriched regulators across all 100 cells and ranked these enriched regulators according to their frequency (Table S3). By comparing the top 30 cell-enriched regulators for each cell type, we identified 39 cell-enriched regulators. Performing the cluster based on the network outdegrees of these 39 cell-enriched regulators from four cell types, we identified cell type-specific regulators and shared regulators (Figure 5C). Notebly, 35 of 39 regulators showed statistically significant differences among four cell types (RPC1, RPC2, PRC3 and MG) in terms of network degree, as indicated by red asterisks in Figure 5C. Interestingly, 33 of these 35 regulators showed significant differences among four cell types, not only in network degree, but also in expression levels (Figure 5D). In particular, the cell-enriched regulators Nr2e3 and Nrl had significantly higher network degrees in MGs than in RPCs, with their expression levels also upregulated in MGs and downregulated in RPCs (Figure 5C and 5D). The Spearman’s correlation coefficients between network degrees and expression levels of the regulators Nr2e3 and Nrl were 0.39 and 0.40, respectively. According to the scATAC-seq data, the regulatory activities of Nr2e3 and Nrl were significantly higher in MGs than in RPCs (Figure S4B). Another regulator Zfp36l1 had a significantly higher network degree in MGs than in RPCs, but its expression level was downregulated in MGs and upregulated in RPCs (Figure 5C and 5D). The Spearman’s correlation coefficient between network degree and expression level of the regulator Zfp36l1 was −0.28. This observation could be due to the fact that Zfp36l1 was negatively regulated by other transcription factors in MGs but not in RPCs (Figure S4C). We also found that Yap1, Dpf2, Hmx1, Pax6, Hmga2, Sox2, Hes1, and Meis1 had higher network degrees in RPCs than in MGs. Accordingly, the expression levels of these factors were upregulated in RPCs and downregulated in MGs. The Spearman’s correlation coefficients between network degrees and expression levels were 0.11, 0.15, 0.11, 0.31, 0.12, 0.15, 0.14 and 0.15, respectively. Additionally, two regulators Chd2 and Peg3 showed network-based cell type specificity, but did not exhibit cell type specificity in terms of gene expression and regulatory activity.

## DISCUSSION

In this study, we developed a novel algorithm, ScReNI, which inferred cell-specific regulatory networks by integrating scRNA-seq and scATAC-seq data. ScReNI employed the nearest neighbors algorithm to identify the neighbors of each cell. It then utilized a random forest with regression trees to model both gene expression and chromatin accessibility, establishing the regulatory relationships among genes. Finally, the random forest was run on each cell and its neighbors to infer cell-specific regulatory networks. Using regulatory relationships detected by ChIP-seq data as the ground truth, we demonstrated that ScReNI outperformed existing methods in predicting cell-specific networks. Moreover, single-cell clustering analysis based on the gene degrees of cell-specific networks also indicated that ScReNI excelled in distinguishing different cell types. Lastly, a unique function in ScReNI was developed to identify cell-enriched regulators, offering new insights into cell-specific regulatory mechanisms.

In current single-cell genomics studies, the integration of scRNA-seq and scATAC-seq data is a challenge due to the technical bath effect. Existing methods for data integration include anchoring methods, transfer learning methods, manifold alignment, matrix factorization and neural network methods (22, 30–33). The advantage of data integrating is that it improves the accuracy of inferring gene regulatory relationships by combining information on gene expression and chromatin accessibility. In our study, we considered the random forest algorithm and developed a method capable of exploiting the synergistic potential of single-cell multi-omics data to construct a cell-specific regulatory network model. This model not only captures the complex interactions between gene expression and chromatin accessibility, but also elucidates potential regulatory relationships at the single-cell level. Our approach provides a robust framework for unraveling the regulatory landscape within individual cells, enabling a deeper understanding of cellular heterogeneity. Furthermore, depending on the characteristics of the dataset, other models, such as neural networks, have the potential to provide better performance in inferring regulatory networks. Future research will include a more comprehensive and theoretically grounded investigation of the different approaches to elucidate their comparative effectiveness in inferring regulatory networks.

When using both gene expression and chromatin accessibility information to construct cell-specific regulatory networks, the key step is to establish the relationships between genes and peaks. The accuracy of predicting these gene-peak relationships can be influenced by factors such as the number of cells analyzed, the sparsity of single-cell sequencing data, and the comprehensiveness of motif databases. In practical applications, these relationships can be predefined, leveraging existing knowledge or biological insights to streamline the analysis process. This capability empowers researchers to focus on the regulatory elements most pertinent to their study, effectively filtering out noise and bolstering the interpretability of the inferred networks.

Despite existing methods, such as SCENIC+ and IReNA (12, 13), showing promising performance in determining key regulators in cell-type-specific networks, it remains a challenge to identify statistically enriched regulators at the single-cell level. Here, we demonstrated that ScReNI could enrich regulators within each cell-specific network. For instance, ScReNI effectively pinpointed transcription factors that acted as regulatory factors for specific cell types by identifying cell-enriched regulators.

These transcription factors, including Nrl and Nr2e3, are notable not only for their regulatory activity but also for their distinctive expression profiles. By identifying cell-enriched regulators, we can unravel the intricate regulatory relationships between transcription factors and target genes, facilitating a more targeted investigation of the underlying regulatory biology.

The cell-enriched regulators Nrl and Nr2e3 had higher network degrees in MGs than in RPCs, with their expression levels upregulated in MGs and downregulated in RPCs. This is consistent with the previous report that Nrl could activate Nr2e3 to suppress photoreceptor development (34). In addition, the enriched regulators Yap1, Dpf2, Hmx1, Pax6, Hmga2, Sox2, Hes1, and Meis1, whose network degrees were higher in RPCs than in MGs, showed a similar pattern of upregulation in RPCs and downregulation in MGs in terms of their expression levels. All of these regulators, except Dpf, have been reported to regulate retinal progenitor cells (35–38). Notably, Chd2 and Peg3 showed a significant difference in network degree among four cell types, but not in expression level or regulatory activity, consistent with the previous concept of ‘dark’ genes (14). The study demonstrated that ScReNI can reliably infer cell-specific regulatory networks and efficiently identify cell-enriched regulators, which has important implications for the regulatory study of diverse biological processes, such as tissue development.

## DATA AVAILABILITY

The unpaired scATAC-seq and scRNA-seq datasets utilized in this study, which pertain to retinal development, can be accessed at GEO accession number GSE181251 (20). Additionally, the scRNA-seq data has been retrieved from a publicly available GitHub repository (https://github.com/gofflab/developing_mouse_retina_scRNASeq). Paired scATAC-seq and scRNA-seq dataset of PBMC from a healthy donor is available on the 10X genomic official website (https://www.10xgenomics.com/datasets/pbmc-from-a-healthy-donor-no-cell-sorting-10-k-1-standard-2-0-0).

## CODE AVAILABILITY

The R package of ScReNI is available on GitHub (https://github.com/Xuxl2020/ScReNI).

## ACKNOWLEDGEMENTS

This study was supported by the National Natural Science Foundation of China [No. T2222003, 32170849], the Ministry of Science and Technology of China [No. 2022YFA1105400], and the Guangdong Province Science and Technology Program [No. 2023B1212060050, 2020B1212060052].

## AUTHOR CONTRIBUTIONS

JW and YL conceived the project and supervised the research. XX, YL, and MT developed the major functions of ScReNI. JLW and XW collected data to check the functions of ScReNI. XX and YL revised the manual of ScReNI. XX and JW drafted the manuscript. All authors contributed to the manuscript revision.

