## Supplementary Information for "ScReNI: single-cell regulatory network inference through integrating scRNA-seq and scATAC-seq data"

**Figure S1.** Illustration of single-cell network inference methods. (A) Workflow of ScReNI. (B) Schematics of the association of gene expression from scRNA-seq and peak accessibility from scATAC-seq.

**Figure S2.** Analysis of top regulation pairs in the retina using the top 2000 highly variable genes. (A) Precision and recall of single-cell regulatory relationships using the same number of regulation pairs in CSN. (B) Precision and recall of single-cell regulatory relationships using different top numbers of regulation pairs.

**Figure S3.** Analysis of the top regulation pairs in peripheral blood mononuclear cells. (A) Precision and recall of single-cell regulatory relationships for 500 highly variable genes using different top numbers of regulation pairs. (B) Precision and recall of single-cell regulatory relationships for 2000 highly variable genes using the same number of regulation pairs in CSN. (C) Precision and recall of single-cell regulatory relationships for 2000 highly variable genes using different top numbers of regulation pairs.

**Figure S4.** Enrichment analysis of the top highly variable genes in retinal development. (A) Enriched functions for genes across six modules. (B) Activities of cell-enriched regulators. P < 0.05 among four cell types was marked with a red star. (C) Average number of genes regulating Zfp36l1 negatively and positively in MGs and RPCs.

**Table S1.** Comparison of network-based cell clustering in retinal development. The maximum ARI values are highlighted in bold.

**Table S2.** Comparison of network-based cell clustering in peripheral blood mononuclear cells. The maximum ARI values are highlighted in bold.

**Table S3.** The list of frequencies of cell-enriched regulators in MG, RPC1, RPC2 and RPC3.

**Figure S1.** Illustration of single-cell network inference methods. (A) Workflow of ScReNI. (B) Schematics of the association of gene expression from scRNA-seq and peak accessibility from scATAC-seq.

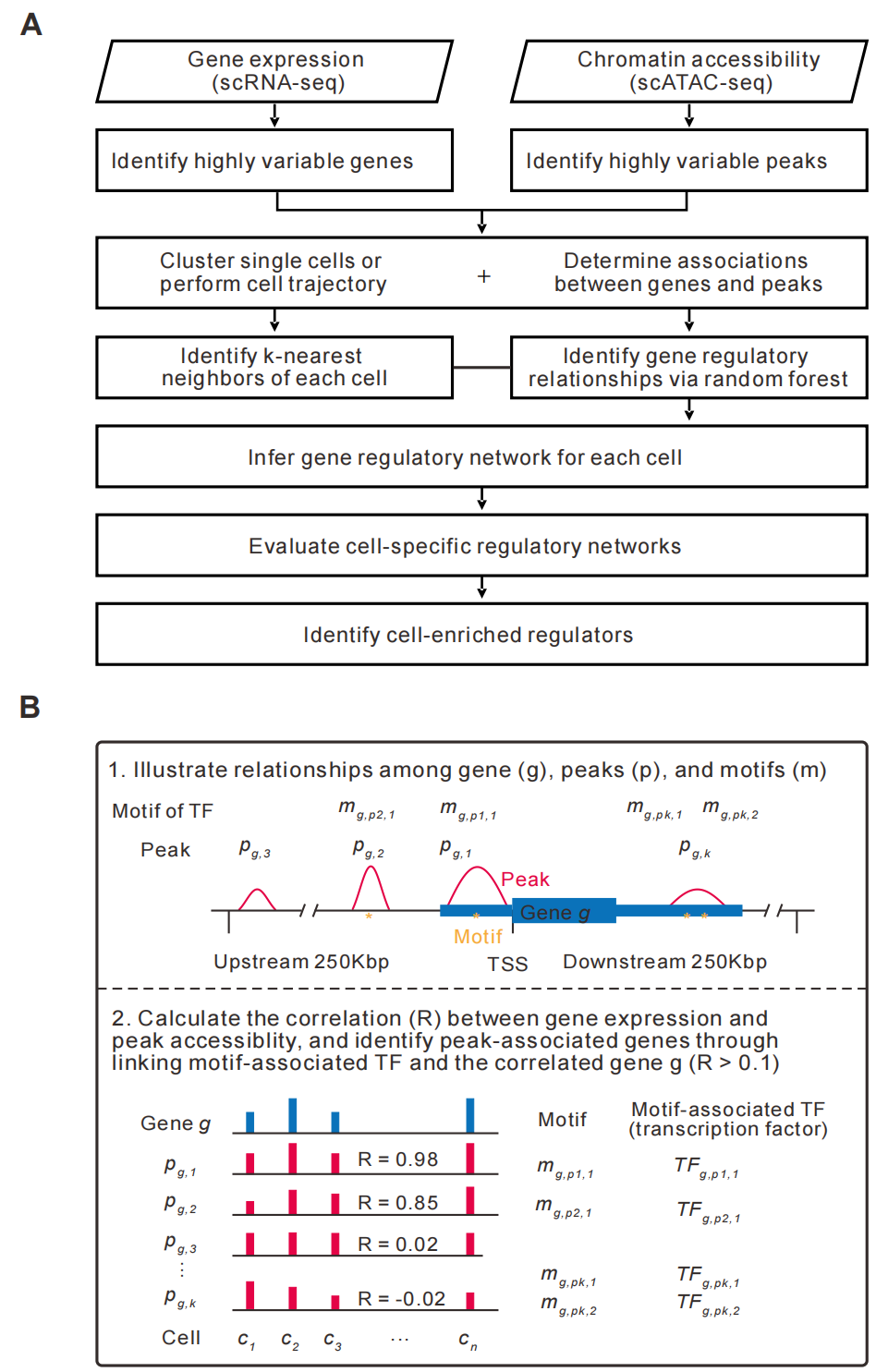

**Figure S2.** Analysis of top regulation pairs in the retina using the top 2000 highly variable genes. (A) Precision and recall of single-cell regulatory relationships using the same number of regulation pairs in CSN. (B) Precision and recall of single-cell regulatory relationships using different top numbers of regulation pairs.

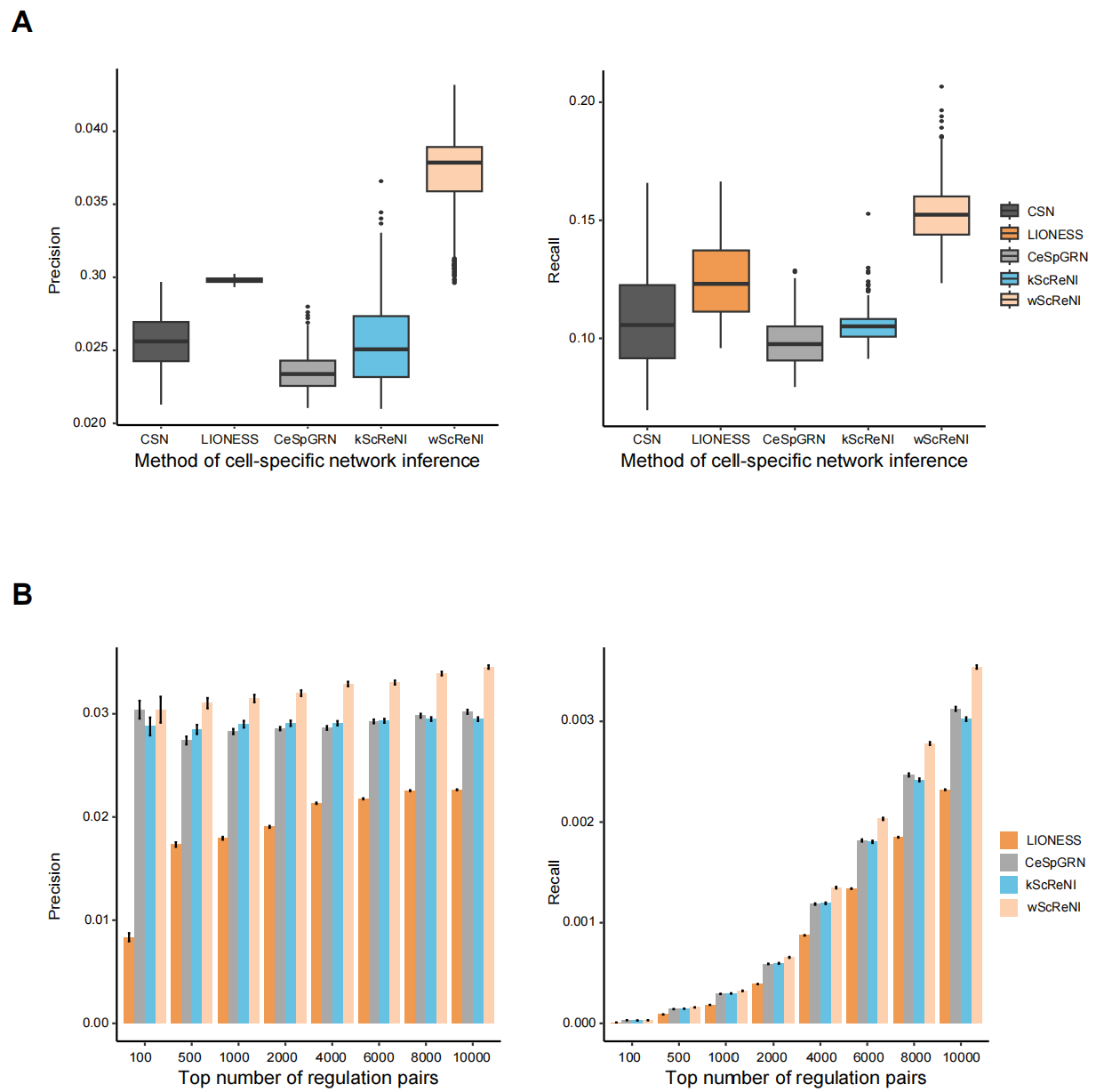

**Figure S3.** Analysis of the top regulation pairs in peripheral blood mononuclear cells. (A) Precision and recall of single-cell regulatory relationships for 500 highly variable genes using different top numbers of regulation pairs. (B) Precision and recall of single-cell regulatory relationships for 2000 highly variable genes using the same number of regulation pairs in CSN. (C) Precision and recall of single-cell regulatory relationships for 2000 highly variable genes using different top numbers of regulation pairs.

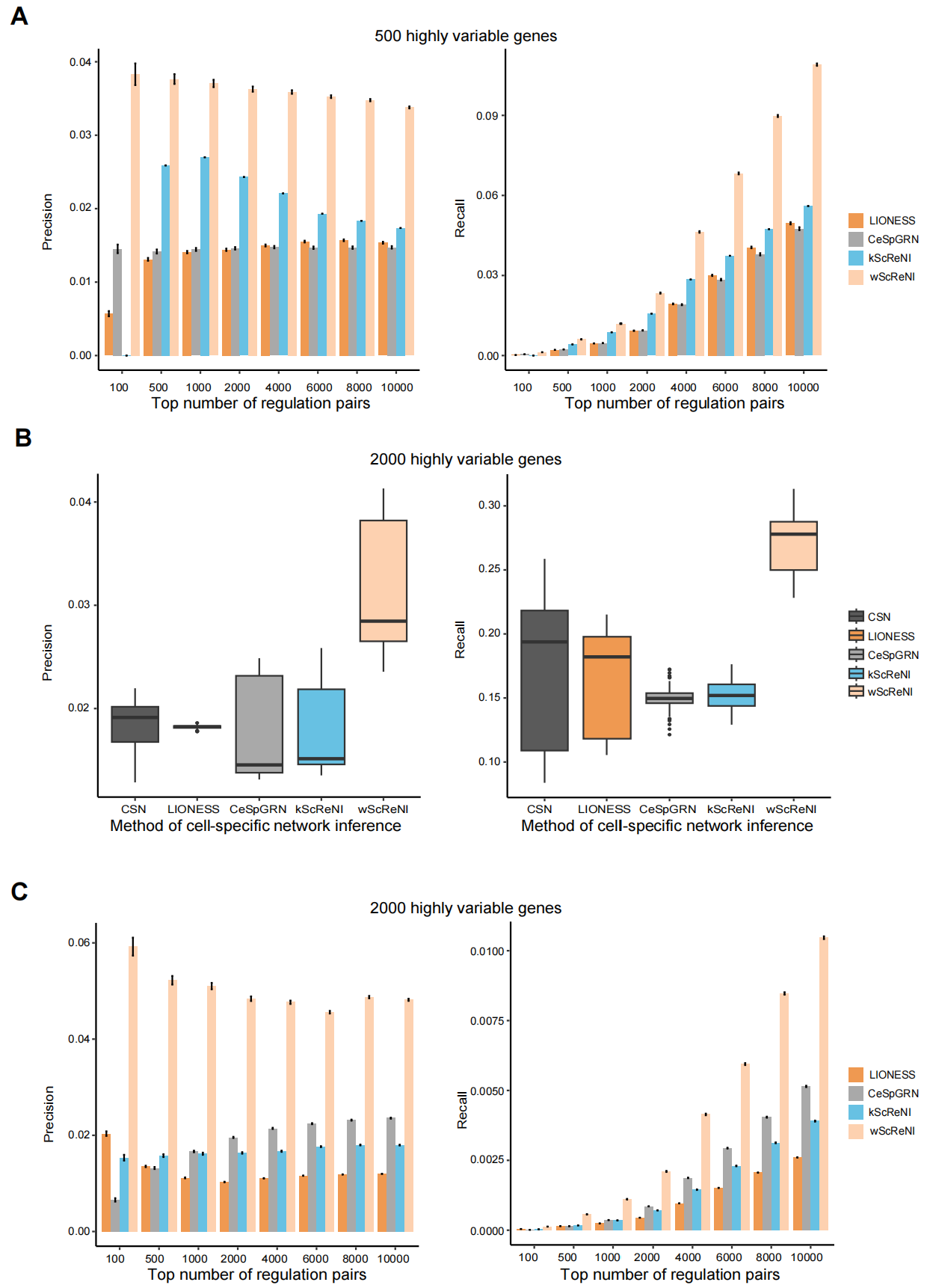

**Figure S4.** Enrichment analysis of the top highly variable genes in retinal development. (A) Enriched functions for genes across six modules. (B) Activities of cell-enriched regulators. P < 0.05 among four cell types was marked with a red star. (C) Average number of genes regulating Zfp36l1 negatively and positively in MGs and RPCs.

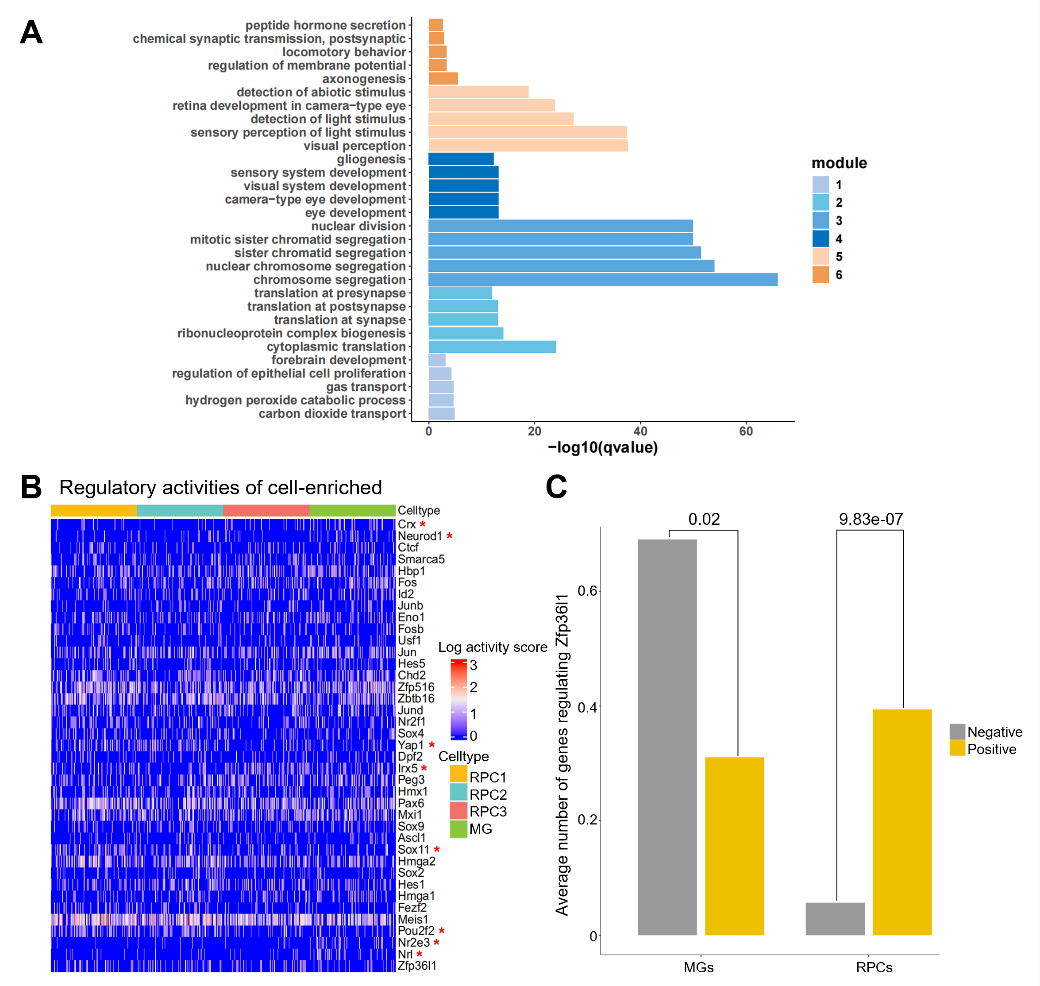

**Table S1.** Comparison of network-based cell clustering in retinal development. The maximum ARI values are highlighted in bold.

| # of genes | Regulation pairs | Types of clustering | CSN | LIONESS | kScReNI | CeSpGRN | wScReNI |
| --- | --- | --- | --- | --- | --- | --- | --- |
| 500 | Top 500 | umap.ARI | 0.488 | 0.001 | 0.517 | 0.513 | **0.642** |
|  |  | hclust.ARI | 0.226 | 0.002 | 0.388 | 0.238 | **0.481** |
|  | Top 800 | umap.ARI | 0.488 | 0.001 | 0.472 | 0.522 | **0.637** |
|  |  | hclust.ARI | 0.226 | 0.002 | 0.244 | 0.238 | **0.466** |
|  | Top 1000 | umap.ARI | 0.488 | 0.001 | 0.521 | 0.453 | **0.647** |
|  |  | hclust.ARI | 0.226 | 0.001 | **0.244** | 0.238 | 0.236 |
|  | Top 3000 | umap.ARI | 0.488 | 0.001 | 0.473 | 0.524 | **0.595** |
|  |  | hclust.ARI | 0.226 | 0.002 | **0.244** | 0.238 | 0.237 |
| 2000 | Top 500 | umap.ARI | 0.411 | 0.001 | 0.483 | 0.469 | **0.698** |
|  |  | hclust.ARI | 0.263 | 0.003 | 0.235 | **0.534** | 0.519 |
|  | Top 800 | umap.ARI | 0.411 | 0.001 | 0.479 | 0.56 | **0.701** |
|  |  | hclust.ARI | 0.263 | 0.004 | 0.245 | 0.521 | **0.525** |
|  | Top 1000 | umap.ARI | 0.411 | 0.001 | 0.479 | 0.58 | **0.715** |
|  |  | hclust.ARI | 0.263 | 0.003 | 0.247 | **0.552** | 0.512 |
|  | Top 3000 | umap.ARI | 0.411 | 0.001 | 0.469 | 0.576 | **0.716** |
|  |  | hclust.ARI | 0.263 | 0.002 | 0.273 | 0.513 | **0.521** |
|  | Top CSN | umap.ARI | 0.411 | 0.144 | 0.522 | 0.617 | **0.636** |
|  |  | hclust.ARI | 0.263 | 0.169 | **0.313** | 0.249 | 0.296 |

**Table S2.** Comparison of network-based cell clustering in peripheral blood mononuclear cells. The maximum ARI values are highlighted in bold.

| # of genes | Regulation pairs | Types of clustering | CSN | LIONESS | kScReNI | CeSpGRN | wScReNI |
| --- | --- | --- | --- | --- | --- | --- | --- |
| 500 | Top 500 | umap.ARI | 0.609 | 0.003 | 0.744 | 0.727 | **0.867** |
|  |  | hclust.ARI | 0.300 | -0.006 | 0.708 | 0.665 | **0.758** |
|  | Top 800 | umap.ARI | 0.609 | 0.001 | 0.744 | 0.773 | **0.863** |
|  |  | hclust.ARI | 0.299 | 0.005 | 0.696 | 0.655 | **0.796** |
|  | Top 1000 | umap.ARI | 0.609 | 0.001 | 0.749 | 0.768 | **0.852** |
|  |  | hclust.ARI | 0.299 | 0.003 | 0.735 | 0.737 | **0.738** |
|  | Top 3000 | umap.ARI | 0.609 | 0.001 | 0.759 | 0.782 | **0.835** |
|  |  | hclust.ARI | 0.299 | 0.002 | 0.725 | 0.753 | **0.778** |
| 2000 | Top 500 | umap.ARI | 0.674 | 0.001 | 0.751 | 0.8 | **0.892** |
|  |  | hclust.ARI | 0.311 | 0.002 | 0.496 | 0.745 | **0.734** |
|  | Top 800 | umap.ARI | 0.674 | 0.001 | 0.762 | 0.763 | **0.893** |
|  |  | hclust.ARI | 0.311 | 0.001 | 0.502 | 0.731 | **0.857** |
|  | Top 1000 | umap.ARI | 0.674 | 0.001 | 0.762 | 0.763 | **0.887** |
|  |  | hclust.ARI | 0.311 | 0.005 | 0.563 | 0.731 | **0.819** |
|  | Top 3000 | umap.ARI | 0.674 | 0.001 | 0.765 | 0.766 | **0.876** |
|  |  | hclust.ARI | 0.311 | 0.001 | 0.57 | 0.677 | **0.72** |
|  | Top CSN | umap.ARI | 0.674 | 0.285 | 0.764 | 0.783 | **0.866** |
|  |  | hclust.ARI | 0.311 | 0.268 | 0.601 | 0.542 | **0.722** |

**Table S3.** The list of frequencies of cell-enriched regulators in MG, RPC1, RPC2 and RPC3.

| Index | Cell type | Enriched regulator | Frequency of regulator present in 100 cells | Index | Cell type | Enriched regulator | Frequency of  regulator present in 100 cells |
| --- | --- | --- | --- | --- | --- | --- | --- |
| 1 | MG | Mxi1 | 74 | 37 | MG | Crx | 48 |
| 2 | MG | Nr2e3 | 74 | 38 | MG | Jund | 48 |
| 3 | MG | Hmga2 | 70 | 39 | MG | Hbp1 | 47 |
| 4 | MG | Zfp516 | 69 | 40 | MG | Maf | 47 |
| 5 | MG | Meis1 | 68 | 41 | MG | Sox8 | 46 |
| 6 | MG | Smarca5 | 68 | 42 | MG | Yap1 | 46 |
| 7 | MG | Zfp36l1 | 68 | 43 | MG | Eno1 | 45 |
| 8 | MG | Pou2f2 | 67 | 44 | MG | Rarb | 45 |
| 9 | MG | Sox11 | 67 | 45 | MG | Junb | 44 |
| 10 | MG | Sox2 | 63 | 46 | MG | Sox5 | 44 |
| 11 | MG | Jun | 62 | 47 | MG | Fosb | 42 |
| 12 | MG | Nr2f1 | 62 | 48 | MG | E2f4 | 39 |
| 13 | MG | Sox4 | 62 | 49 | MG | Myc | 36 |
| 14 | MG | Dpf2 | 61 | 50 | MG | Ctcf | 35 |
| 15 | MG | Fezf2 | 61 | 51 | MG | E2f3 | 35 |
| 16 | MG | Mafb | 61 | 52 | MG | Foxp1 | 34 |
| 17 | MG | Pax6 | 60 | 53 | MG | Egr1 | 33 |
| 18 | MG | Peg3 | 60 | 54 | MG | Klf13 | 31 |
| 19 | MG | Ppard | 59 | 55 | MG | Nfix | 31 |
| 20 | MG | Pou2f1 | 58 | 56 | MG | Thrb | 30 |
| 21 | MG | Sox9 | 58 | 57 | MG | Bach1 | 29 |
| 22 | MG | Usf1 | 57 | 58 | MG | Zfp236 | 29 |
| 23 | MG | Fos | 56 | 59 | MG | Nfib | 28 |
| 24 | MG | Neurod1 | 55 | 60 | MG | Id2 | 27 |
| 25 | MG | Pou3f2 | 55 | 61 | MG | Meis2 | 26 |
| 26 | MG | Hmga1 | 54 | 62 | MG | Thra | 26 |
| 27 | MG | Chd2 | 53 | 63 | MG | Zscan26 | 24 |
| 28 | MG | Klf9 | 53 | 64 | MG | Creb1 | 23 |
| 29 | MG | Nfic | 53 | 65 | MG | Klf2 | 23 |
| 30 | MG | Hes5 | 52 | 66 | MG | Pknox1 | 23 |
| 31 | MG | Nr2f2 | 52 | 67 | MG | Nr2f6 | 21 |
| 32 | MG | Nrl | 52 | 68 | MG | Prdm1 | 20 |
| 33 | MG | Hes1 | 51 | 69 | MG | Hnrnpul1 | 19 |
| 34 | MG | Irx5 | 51 | 70 | MG | Mxd1 | 18 |
| 35 | MG | Otx2 | 51 | 71 | MG | Rfx3 | 18 |
| 36 | MG | Ascl1 | 50 | 72 | MG | Vsx2 | 18 |
| 73 | MG | Gtf3c2 | 17 | 117 | MG | Tgif1 | 2 |
| 74 | MG | Nfia | 17 | 118 | MG | Zfp251 | 2 |
| 75 | MG | Srf | 17 | 119 | MG | Zfp667 | 2 |
| 76 | MG | Dbp | 16 | 120 | MG | Bhlhe23 | 1 |
| 77 | MG | Zxdc | 16 | 121 | MG | Gli1 | 1 |
| 78 | MG | Zfp266 | 15 | 122 | MG | Hic2 | 1 |
| 79 | MG | Zfp746 | 15 | 123 | MG | Lmo2 | 1 |
| 80 | MG | Foxn3 | 14 | 124 | MG | Mybl1 | 1 |
| 81 | MG | Insm1 | 14 | 125 | MG | Neurog2 | 1 |
| 82 | MG | Nfatc3 | 14 | 126 | MG | Nhlh2 | 1 |
| 83 | MG | Xbp1 | 14 | 127 | MG | Olig2 | 1 |
| 84 | MG | Terf1 | 12 | 128 | MG | Otx1 | 1 |
| 85 | MG | Zfp568 | 12 | 129 | MG | Prox1 | 1 |
| 86 | MG | Tfap2b | 11 | 130 | MG | Zbtb39 | 1 |
| 87 | MG | Gsc2 | 10 | 131 | MG | Zfp711 | 1 |
| 88 | MG | Mxd3 | 10 | 1 | RPC1 | Sox4 | 72 |
| 89 | MG | Atf6 | 8 | 2 | RPC1 | Pou2f2 | 71 |
| 90 | MG | Dnttip1 | 8 | 3 | RPC1 | Smarca5 | 66 |
| 91 | MG | Rbpj | 8 | 4 | RPC1 | Sox2 | 65 |
| 92 | MG | Stat1 | 8 | 5 | RPC1 | Usf1 | 64 |
| 93 | MG | Zic1 | 8 | 6 | RPC1 | Dpf2 | 63 |
| 94 | MG | Nr2c1 | 6 | 7 | RPC1 | Irx5 | 63 |
| 95 | MG | Nr2e1 | 6 | 8 | RPC1 | Nr2f1 | 62 |
| 96 | MG | Tfe3 | 6 | 9 | RPC1 | Zfp36l1 | 62 |
| 97 | MG | Zbtb48 | 6 | 10 | RPC1 | Hmga2 | 61 |
| 98 | MG | Hey1 | 5 | 11 | RPC1 | Meis1 | 61 |
| 99 | MG | Lhx4 | 5 | 12 | RPC1 | Fezf2 | 60 |
| 100 | MG | Rest | 5 | 13 | RPC1 | Jun | 60 |
| 101 | MG | Vsx1 | 5 | 14 | RPC1 | Mxi1 | 60 |
| 102 | MG | Foxn4 | 4 | 15 | RPC1 | Sox11 | 60 |
| 103 | MG | Zfhx3 | 4 | 16 | RPC1 | Foxp1 | 59 |
| 104 | MG | Arid5b | 3 | 17 | RPC1 | Zfp516 | 59 |
| 105 | MG | Atf3 | 3 | 18 | RPC1 | Peg3 | 57 |
| 106 | MG | Ebf1 | 3 | 19 | RPC1 | Chd2 | 56 |
| 107 | MG | Foxg1 | 3 | 20 | RPC1 | Pax6 | 55 |
| 108 | MG | Foxp2 | 3 | 21 | RPC1 | Fos | 54 |
| 109 | MG | Gmeb1 | 3 | 22 | RPC1 | Pou2f1 | 54 |
| 110 | MG | Hmx1 | 3 | 23 | RPC1 | Klf13 | 53 |
| 111 | MG | Id4 | 3 | 24 | RPC1 | Maf | 53 |
| 112 | MG | Zbtb16 | 3 | 25 | RPC1 | Mafb | 53 |
| 113 | MG | Isl1 | 2 | 26 | RPC1 | Nr2e3 | 52 |
| 114 | MG | Mecom | 2 | 27 | RPC1 | Hmga1 | 48 |
| 115 | MG | Rnf141 | 2 | 28 | RPC1 | Nr2f2 | 48 |
| 116 | MG | Tbx3 | 2 | 29 | RPC1 | Jund | 47 |
| 30 | RPC1 | Sox9 | 47 | 74 | RPC1 | Pknox1 | 19 |
| 31 | RPC1 | Yap1 | 47 | 75 | RPC1 | Bach1 | 18 |
| 32 | RPC1 | Myc | 46 | 76 | RPC1 | E2f3 | 18 |
| 33 | RPC1 | Hbp1 | 44 | 77 | RPC1 | Vsx2 | 18 |
| 34 | RPC1 | Rarb | 43 | 78 | RPC1 | Foxd1 | 17 |
| 35 | RPC1 | Eno1 | 41 | 79 | RPC1 | Klf9 | 17 |
| 36 | RPC1 | Hes1 | 41 | 80 | RPC1 | Meis2 | 17 |
| 37 | RPC1 | Hes5 | 41 | 81 | RPC1 | Mxd3 | 17 |
| 38 | RPC1 | Sox8 | 41 | 82 | RPC1 | Zfp266 | 17 |
| 39 | RPC1 | E2f4 | 39 | 83 | RPC1 | Zfp711 | 16 |
| 40 | RPC1 | Tbx5 | 38 | 84 | RPC1 | Foxg1 | 15 |
| 41 | RPC1 | Creb1 | 36 | 85 | RPC1 | Bcl11b | 13 |
| 42 | RPC1 | Ctcf | 36 | 86 | RPC1 | Zbtb48 | 13 |
| 43 | RPC1 | Zic1 | 36 | 87 | RPC1 | Atf3 | 12 |
| 44 | RPC1 | Ppard | 35 | 88 | RPC1 | Foxp2 | 12 |
| 45 | RPC1 | Ascl1 | 34 | 89 | RPC1 | Klf2 | 12 |
| 46 | RPC1 | Pou3f2 | 33 | 90 | RPC1 | Stat1 | 12 |
| 47 | RPC1 | Rfx3 | 33 | 91 | RPC1 | Zscan26 | 11 |
| 48 | RPC1 | Thra | 33 | 92 | RPC1 | Hinfp | 10 |
| 49 | RPC1 | Hmx1 | 32 | 93 | RPC1 | Srf | 10 |
| 50 | RPC1 | Junb | 31 | 94 | RPC1 | Xbp1 | 10 |
| 51 | RPC1 | Nrl | 31 | 95 | RPC1 | Zxdc | 10 |
| 52 | RPC1 | Rest | 30 | 96 | RPC1 | Insm1 | 9 |
| 53 | RPC1 | Id2 | 29 | 97 | RPC1 | Foxn4 | 8 |
| 54 | RPC1 | Egr1 | 28 | 98 | RPC1 | Mxd1 | 8 |
| 55 | RPC1 | Otx1 | 28 | 99 | RPC1 | Nfic | 8 |
| 56 | RPC1 | Hnrnpul1 | 27 | 100 | RPC1 | Zfhx3 | 8 |
| 57 | RPC1 | Nfib | 27 | 101 | RPC1 | Dnttip1 | 7 |
| 58 | RPC1 | Sox5 | 25 | 102 | RPC1 | Rnf141 | 7 |
| 59 | RPC1 | Mecom | 24 | 103 | RPC1 | Terf1 | 7 |
| 60 | RPC1 | Otx2 | 24 | 104 | RPC1 | Zfp568 | 7 |
| 61 | RPC1 | Fosb | 23 | 105 | RPC1 | Hic2 | 6 |
| 62 | RPC1 | Rbpj | 23 | 106 | RPC1 | Nr2c1 | 6 |
| 63 | RPC1 | Tbx3 | 23 | 107 | RPC1 | Zfp251 | 6 |
| 64 | RPC1 | Dbp | 22 | 108 | RPC1 | Zfp746 | 6 |
| 65 | RPC1 | Gtf3c2 | 22 | 109 | RPC1 | Gsc2 | 5 |
| 66 | RPC1 | Zfp667 | 22 | 110 | RPC1 | Mybl1 | 5 |
| 67 | RPC1 | Nr2e1 | 21 | 111 | RPC1 | Ebf1 | 4 |
| 68 | RPC1 | Tgif1 | 21 | 112 | RPC1 | Isl1 | 4 |
| 69 | RPC1 | Foxn3 | 20 | 113 | RPC1 | Pitx3 | 4 |
| 70 | RPC1 | Zbtb16 | 20 | 114 | RPC1 | Hey1 | 3 |
| 71 | RPC1 | Zfp236 | 20 | 115 | RPC1 | Neurod1 | 3 |
| 72 | RPC1 | Nfatc3 | 19 | 116 | RPC1 | Atf6 | 2 |
| 73 | RPC1 | Nr2f6 | 19 | 117 | RPC1 | Gmeb1 | 2 |
| 118 | RPC1 | Id4 | 2 | 39 | RPC2 | Yap1 | 42 |
| 119 | RPC1 | Irf8 | 1 | 40 | RPC2 | Pou3f2 | 40 |
| 120 | RPC1 | Lhx4 | 1 | 41 | RPC2 | Rest | 40 |
| 121 | RPC1 | Msx1 | 1 | 42 | RPC2 | Eno1 | 39 |
| 122 | RPC1 | Neurog2 | 1 | 43 | RPC2 | Myc | 39 |
| 123 | RPC1 | Prox1 | 1 | 44 | RPC2 | Klf13 | 36 |
| 1 | RPC2 | Usf1 | 68 | 45 | RPC2 | Sox5 | 36 |
| 2 | RPC2 | Pou2f2 | 67 | 46 | RPC2 | Rarb | 34 |
| 3 | RPC2 | Zfp516 | 67 | 47 | RPC2 | Id2 | 32 |
| 4 | RPC2 | Smarca5 | 64 | 48 | RPC2 | Nrl | 32 |
| 5 | RPC2 | Hmga2 | 63 | 49 | RPC2 | Tbx3 | 32 |
| 6 | RPC2 | Jun | 63 | 50 | RPC2 | Zic1 | 32 |
| 7 | RPC2 | Fos | 62 | 51 | RPC2 | Otx2 | 30 |
| 8 | RPC2 | Maf | 62 | 52 | RPC2 | Creb1 | 29 |
| 9 | RPC2 | Peg3 | 62 | 53 | RPC2 | Junb | 29 |
| 10 | RPC2 | Dpf2 | 60 | 54 | RPC2 | Mecom | 29 |
| 11 | RPC2 | Hes5 | 59 | 55 | RPC2 | Klf2 | 28 |
| 12 | RPC2 | Mxi1 | 59 | 56 | RPC2 | Nfib | 28 |
| 13 | RPC2 | Sox11 | 59 | 57 | RPC2 | Foxn3 | 27 |
| 14 | RPC2 | Irx5 | 58 | 58 | RPC2 | Nr2e1 | 27 |
| 15 | RPC2 | Nr2e3 | 58 | 59 | RPC2 | Nr2f6 | 26 |
| 16 | RPC2 | Pou2f1 | 58 | 60 | RPC2 | Pknox1 | 26 |
| 17 | RPC2 | Sox9 | 58 | 61 | RPC2 | Gtf3c2 | 25 |
| 18 | RPC2 | Fezf2 | 57 | 62 | RPC2 | Rbpj | 25 |
| 19 | RPC2 | Zfp36l1 | 57 | 63 | RPC2 | Bach1 | 24 |
| 20 | RPC2 | Nr2f1 | 56 | 64 | RPC2 | E2f4 | 23 |
| 21 | RPC2 | Meis1 | 55 | 65 | RPC2 | Foxn4 | 23 |
| 22 | RPC2 | Hes1 | 54 | 66 | RPC2 | Mybl1 | 23 |
| 23 | RPC2 | Jund | 54 | 67 | RPC2 | Otx1 | 23 |
| 24 | RPC2 | Nr2f2 | 53 | 68 | RPC2 | Thra | 23 |
| 25 | RPC2 | Pax6 | 52 | 69 | RPC2 | Zfp711 | 23 |
| 26 | RPC2 | Foxp1 | 49 | 70 | RPC2 | Hmx1 | 22 |
| 27 | RPC2 | Hmga1 | 49 | 71 | RPC2 | Hnrnpul1 | 22 |
| 28 | RPC2 | Chd2 | 48 | 72 | RPC2 | Klf9 | 22 |
| 29 | RPC2 | Sox2 | 48 | 73 | RPC2 | Zbtb16 | 22 |
| 30 | RPC2 | Sox4 | 47 | 74 | RPC2 | Foxg1 | 21 |
| 31 | RPC2 | Mafb | 46 | 75 | RPC2 | Foxd1 | 20 |
| 32 | RPC2 | Ppard | 46 | 76 | RPC2 | Foxp2 | 20 |
| 33 | RPC2 | Ascl1 | 45 | 77 | RPC2 | Meis2 | 19 |
| 34 | RPC2 | Egr1 | 45 | 78 | RPC2 | Neurod1 | 19 |
| 35 | RPC2 | Ctcf | 44 | 79 | RPC2 | Tbx5 | 19 |
| 36 | RPC2 | Hbp1 | 43 | 80 | RPC2 | Zfp568 | 19 |
| 37 | RPC2 | Sox8 | 43 | 81 | RPC2 | Mxd3 | 18 |
| 38 | RPC2 | Fosb | 42 | 82 | RPC2 | Tgif1 | 17 |
| 83 | RPC2 | Neurog2 | 16 | 127 | RPC2 | Prox1 | 2 |
| 84 | RPC2 | Zfp746 | 15 | 128 | RPC2 | Thrb | 2 |
| 85 | RPC2 | Zscan26 | 15 | 129 | RPC2 | Lmo2 | 1 |
| 86 | RPC2 | Dbp | 14 | 130 | RPC2 | Nhlh2 | 1 |
| 87 | RPC2 | Rfx3 | 14 | 131 | RPC2 | Tfap2b | 1 |
| 88 | RPC2 | Vsx2 | 14 | 132 | RPC2 | Tfe3 | 1 |
| 89 | RPC2 | Insm1 | 13 | 1 | RPC3 | Fezf2 | 72 |
| 90 | RPC2 | Pitx3 | 13 | 2 | RPC3 | Hes5 | 72 |
| 91 | RPC2 | Zfp266 | 13 | 3 | RPC3 | Nr2e3 | 72 |
| 92 | RPC2 | E2f3 | 12 | 4 | RPC3 | Nr2f1 | 71 |
| 93 | RPC2 | Id4 | 12 | 5 | RPC3 | Pou2f2 | 70 |
| 94 | RPC2 | Xbp1 | 12 | 6 | RPC3 | Mxi1 | 68 |
| 95 | RPC2 | Hic2 | 11 | 7 | RPC3 | Meis1 | 66 |
| 96 | RPC2 | Srf | 11 | 8 | RPC3 | Zfp516 | 66 |
| 97 | RPC2 | Terf1 | 11 | 9 | RPC3 | Sox2 | 65 |
| 98 | RPC2 | Zfp236 | 11 | 10 | RPC3 | Peg3 | 64 |
| 99 | RPC2 | Mxd1 | 10 | 11 | RPC3 | Dpf2 | 63 |
| 100 | RPC2 | Nfia | 10 | 12 | RPC3 | Hmga2 | 63 |
| 101 | RPC2 | Hey1 | 9 | 13 | RPC3 | Sox8 | 61 |
| 102 | RPC2 | Rnf141 | 9 | 14 | RPC3 | Maf | 59 |
| 103 | RPC2 | Zxdc | 9 | 15 | RPC3 | Smarca5 | 59 |
| 104 | RPC2 | Ebf1 | 8 | 16 | RPC3 | Chd2 | 56 |
| 105 | RPC2 | Gli1 | 8 | 17 | RPC3 | Irx5 | 56 |
| 106 | RPC2 | Nfatc3 | 8 | 18 | RPC3 | Jun | 55 |
| 107 | RPC2 | Hinfp | 7 | 19 | RPC3 | Sox4 | 55 |
| 108 | RPC2 | Nfix | 7 | 20 | RPC3 | Ascl1 | 54 |
| 109 | RPC2 | Zfp667 | 7 | 21 | RPC3 | Sox9 | 54 |
| 110 | RPC2 | Arid5b | 6 | 22 | RPC3 | Usf1 | 53 |
| 111 | RPC2 | Msx1 | 6 | 23 | RPC3 | Yap1 | 53 |
| 112 | RPC2 | Stat1 | 6 | 24 | RPC3 | Zfp36l1 | 53 |
| 113 | RPC2 | Crx | 5 | 25 | RPC3 | Fos | 52 |
| 114 | RPC2 | Zfhx3 | 5 | 26 | RPC3 | Sox11 | 50 |
| 115 | RPC2 | Zbtb48 | 4 | 27 | RPC3 | Foxp1 | 49 |
| 116 | RPC2 | Zfp362 | 4 | 28 | RPC3 | Hmga1 | 49 |
| 117 | RPC2 | Dnttip1 | 3 | 29 | RPC3 | Pou2f1 | 48 |
| 118 | RPC2 | Gmeb1 | 3 | 30 | RPC3 | Pou3f2 | 48 |
| 119 | RPC2 | Irf8 | 3 | 31 | RPC3 | Ppard | 48 |
| 120 | RPC2 | Nfic | 3 | 32 | RPC3 | Jund | 47 |
| 121 | RPC2 | Zbtb39 | 3 | 33 | RPC3 | Nr2f2 | 47 |
| 122 | RPC2 | Zfp251 | 3 | 34 | RPC3 | Egr1 | 45 |
| 123 | RPC2 | Atf3 | 2 | 35 | RPC3 | Rarb | 45 |
| 124 | RPC2 | Atf6 | 2 | 36 | RPC3 | Pax6 | 44 |
| 125 | RPC2 | Isl1 | 2 | 37 | RPC3 | Hes1 | 42 |
| 126 | RPC2 | Nr2c1 | 2 | 38 | RPC3 | Junb | 42 |
| 39 | RPC3 | Sox5 | 42 | 83 | RPC3 | Mxd3 | 18 |
| 40 | RPC3 | Ctcf | 40 | 84 | RPC3 | Vsx2 | 17 |
| 41 | RPC3 | Fosb | 40 | 85 | RPC3 | Dbp | 16 |
| 42 | RPC3 | Nrl | 40 | 86 | RPC3 | Foxp2 | 16 |
| 43 | RPC3 | Eno1 | 39 | 87 | RPC3 | Gtf3c2 | 15 |
| 44 | RPC3 | Mafb | 38 | 88 | RPC3 | Nr2f6 | 15 |
| 45 | RPC3 | Neurod1 | 38 | 89 | RPC3 | Prdm1 | 15 |
| 46 | RPC3 | Klf13 | 36 | 90 | RPC3 | Tbx5 | 14 |
| 47 | RPC3 | Otx2 | 35 | 91 | RPC3 | Zfp746 | 14 |
| 48 | RPC3 | Rest | 35 | 92 | RPC3 | Foxn4 | 13 |
| 49 | RPC3 | Klf9 | 33 | 93 | RPC3 | Irf8 | 13 |
| 50 | RPC3 | Hbp1 | 32 | 94 | RPC3 | Nfatc3 | 13 |
| 51 | RPC3 | Id2 | 30 | 95 | RPC3 | Id4 | 12 |
| 52 | RPC3 | Nfib | 30 | 96 | RPC3 | Hic2 | 11 |
| 53 | RPC3 | Nfix | 28 | 97 | RPC3 | Tfap2b | 11 |
| 54 | RPC3 | Nr2e1 | 28 | 98 | RPC3 | Crx | 10 |
| 55 | RPC3 | Neurog2 | 26 | 99 | RPC3 | Hey1 | 10 |
| 56 | RPC3 | Bach1 | 25 | 100 | RPC3 | Insm1 | 10 |
| 57 | RPC3 | Hnrnpul1 | 25 | 101 | RPC3 | Zfp251 | 10 |
| 58 | RPC3 | Nfic | 25 | 102 | RPC3 | Zfp266 | 10 |
| 59 | RPC3 | Nfia | 24 | 103 | RPC3 | Creb1 | 9 |
| 60 | RPC3 | Pknox1 | 24 | 104 | RPC3 | Rnf141 | 9 |
| 61 | RPC3 | Rfx3 | 24 | 105 | RPC3 | Zbtb16 | 9 |
| 62 | RPC3 | Tbx3 | 24 | 106 | RPC3 | Olig2 | 8 |
| 63 | RPC3 | Thra | 24 | 107 | RPC3 | Terf1 | 8 |
| 64 | RPC3 | Zic1 | 24 | 108 | RPC3 | Thrb | 8 |
| 65 | RPC3 | E2f4 | 23 | 109 | RPC3 | Zfp711 | 8 |
| 66 | RPC3 | Mybl1 | 23 | 110 | RPC3 | Zxdc | 8 |
| 67 | RPC3 | Klf2 | 22 | 111 | RPC3 | Foxd1 | 7 |
| 68 | RPC3 | Gli1 | 21 | 112 | RPC3 | Ptf1a | 7 |
| 69 | RPC3 | Myc | 21 | 113 | RPC3 | Arid5b | 6 |
| 70 | RPC3 | Stat1 | 21 | 114 | RPC3 | Atf6 | 6 |
| 71 | RPC3 | Xbp1 | 21 | 115 | RPC3 | Isl1 | 5 |
| 72 | RPC3 | Foxn3 | 20 | 116 | RPC3 | Lhx4 | 5 |
| 73 | RPC3 | Rbpj | 20 | 117 | RPC3 | Zfp236 | 5 |
| 74 | RPC3 | Atf3 | 19 | 118 | RPC3 | Zfp667 | 5 |
| 75 | RPC3 | Mxd1 | 19 | 119 | RPC3 | Bcl11b | 4 |
| 76 | RPC3 | Srf | 19 | 120 | RPC3 | Dnttip1 | 4 |
| 77 | RPC3 | Tgif1 | 19 | 121 | RPC3 | Hinfp | 4 |
| 78 | RPC3 | Zscan26 | 19 | 122 | RPC3 | Lmo2 | 4 |
| 79 | RPC3 | E2f3 | 18 | 123 | RPC3 | Mecom | 4 |
| 80 | RPC3 | Foxg1 | 18 | 124 | RPC3 | Nr2c1 | 4 |
| 81 | RPC3 | Hmx1 | 18 | 125 | RPC3 | Otx1 | 4 |
| 82 | RPC3 | Meis2 | 18 | 126 | RPC3 | Tfe3 | 4 |
| 127 | RPC3 | Zbtb39 | 4 | 134 | RPC3 | Zbtb48 | 3 |
| 128 | RPC3 | Zfhx3 | 4 | 135 | RPC3 | Zfp568 | 3 |
| 129 | RPC3 | Zfp362 | 4 | 136 | RPC3 | Gsc2 | 2 |
| 130 | RPC3 | Ebf1 | 3 | 137 | RPC3 | Msx1 | 2 |
| 131 | RPC3 | Gmeb1 | 3 | 138 | RPC3 | Prox1 | 2 |
| 132 | RPC3 | Nhlh2 | 3 | 139 | RPC3 | Vsx1 | 1 |
| 133 | RPC3 | Pitx3 | 3 |  |  |  |  |
